# Neuropeptide Y4 receptor activation delays autoimmune diabetes by reprogramming β-cell stress and immune tolerance

**DOI:** 10.64898/2026.07.03.736290

**Authors:** N.A Haq, K.W Toczyska, A Islam, O.E Olaniru, Y Lei, M Hu, M Zhao, R Müller, M.K.M Mirza, N.H.F. Fine, D. J Hodson, S.J Persaud, A. G Beck-Sickinger, JA Pearson, G.A Bewick

## Abstract

Type 1 diabetes (T1D) involves immune-mediated destruction of pancreatic β-cells, yet current disease-modifying therapies mainly target immunity without enhancing β-cell resilience. We show selective neuropeptide Y4 receptor (Y4R) agonism protects β-cells while reshaping islet immunity across T1D models. Multi-modal localisation using cell sorting, qPCR, RNAscope and fluorescent ligand competition demonstrated predominant Y4R expression and functional accessibility on mouse and human β-cells. Selective Y4R agonism was non-toxic and did not impair islet network integrity, Ca²⁺ dynamics, glucose-stimulated insulin secretion or systemic glucose tolerance. Y4R activation conferred cytoprotection against inflammatory cytokines, streptozotocin, lipotoxicity and ER stress, reducing caspase-3/7 activation and β-cell loss whilst sustaining insulin release and promoting proliferation in both mouse and human islets. Bulk RNA-seq revealed a coordinated β-cell resilience programme characterised by reinforced identity and insulin processing, KEAP1-NFE2L2-driven antioxidative and proteostatic activation, and suppression of EIF2 signalling and associated biosynthetic and ER stress pathways. Concurrently, Y4R agonism dampened pathogenic chemokine and cytokine networks, including CXCL10, CCL3/4/7 and IL-6, while preserving IL-2 and Foxp3 signals, thereby limiting CD8⁺ T cell, CD4⁺ T cell and macrophage chemotaxis toward cytokine-stressed islets. Reduced immune-cell recruitment was conserved in a fully human immune-islet system, where Y4R activation significantly attenuated IL-2-activated human PBMC migration and invasion toward cytokine-stressed human islets. In a stringent NY8.3 CD8⁺ T cell adoptive-transfer model, systemic Y4R agonism significantly delayed diabetes onset. These data position Y4R as a β-cell-centric therapeutic target coupling intrinsic resilience with local immune modulation, offering a complementary approach for β-cell preservation in T1D and islet replacement therapies.

**Graphical abstract:** The selective Y4 receptor agonist K22 binds β-cell–enriched NPY4R in mouse and human islets, activates a β-cell resilience programme that preserves insulin secretion under inflammatory and metabolic stress, and simultaneously dampens islet chemokine output, thereby limiting innate and adaptive immune-cell recruitment and delaying autoimmune diabetes onset.

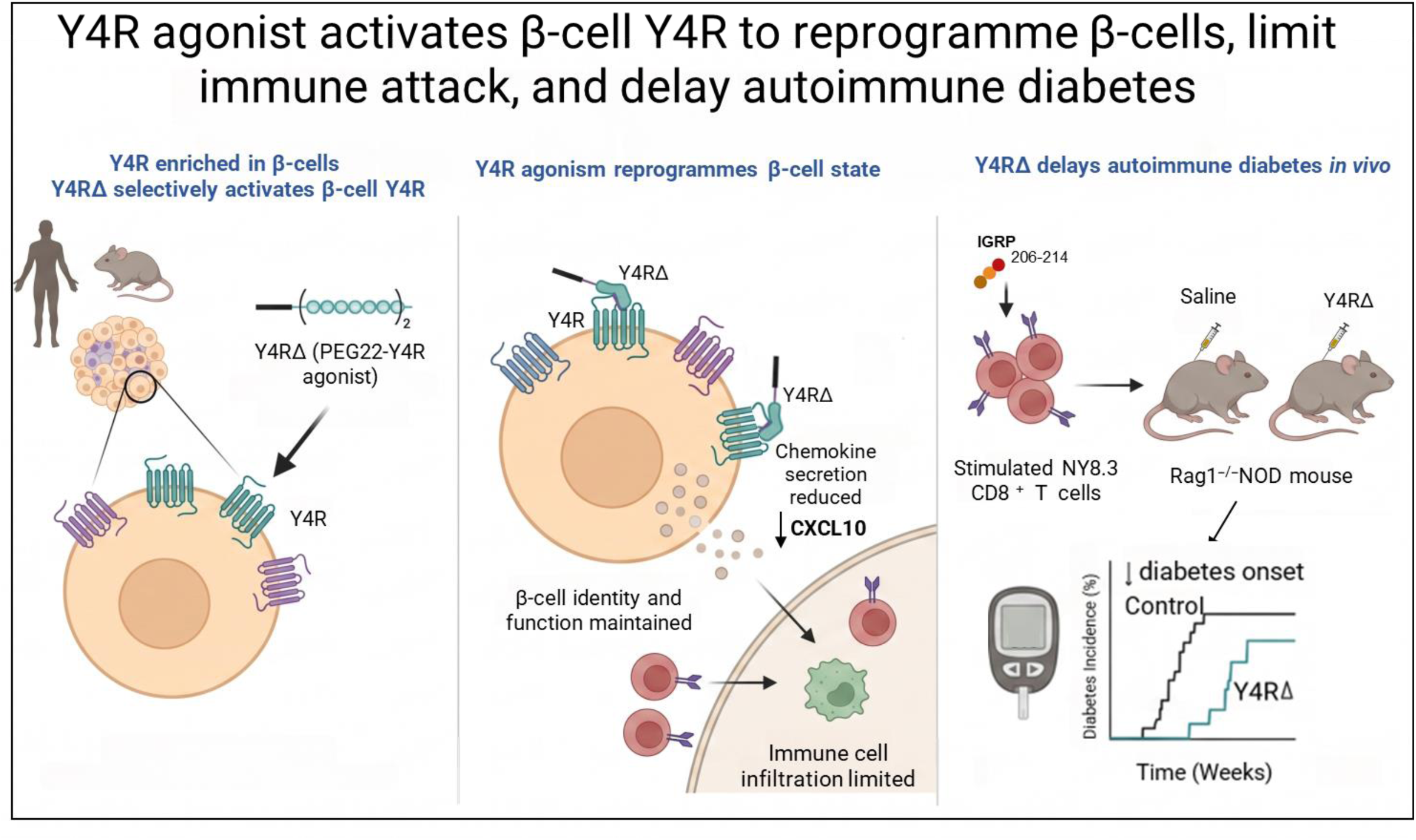

## Introduction

Type 1 diabetes mellitus (T1D) is a chronic autoimmune disease caused by immune-mediated destruction of insulin-producing pancreatic β-cells, resulting in insulin deficiency and dysregulated metabolic homeostasis [1, 2]. T1D affects over 8.4 million people globally, with incidence rising by 2–5% annually, highlighting the urgent need for disease-modifying therapies [3, 4].

Recent approval of teplizumab, an anti-CD3 monoclonal antibody, and promising preclinical results with the JAK inhibitor baricitinib demonstrate the therapeutic potential of immune modulation to delay disease onset. However, these strategies rely on systemic immune modulation (with attendant infection/malignancy risks), which may limit long-term safety and widespread applicability [5, 6]. There remains a critical need for complementary approaches that enhance intrinsic β-cell resilience to immune-mediated damage whilst modulating local immune responses, potentially reducing reliance on chronic immunosuppression.

The neuropeptide Y (NPY) system represents an evolutionarily conserved signalling network with established roles in energy balance, appetite regulation, and stress responses [7, 8]. In peripheral tissues including adipose, gut, and vasculature, NPY and its receptors modulate metabolism, inflammation, and cell survival. Although extensively studied in metabolic regulation, the role of NPY in islet biology remains comparatively understudied [9, 10]. Within islets, NPY is produced by sympathetic nerve terminals and β-cells, and other family members include peptide YY (PYY) produced by α-cells, and pancreatic polypeptide (PP) by PP cells. These peptides signal through class A G-protein coupled receptors (Y1, Y2, Y4, and Y5) each with distinct ligand affinities and downstream signalling.

NPY and PYY1-36 bind *Y1R*, *Y2R*, and *Y5R* receptors with high affinity, whilst PP preferentially activates *Npy4r* [11, 12]. Although *Npy1r* receptor signalling is known to suppress insulin secretion via Gi-mediated inhibition of cyclic AMP, the broader contributions of individual receptor subtypes to islet function under inflammatory stress remain poorly defined [13].

Emerging evidence, including our own, suggests the NPY system may influence β-cell survival and functional plasticity [14–16]. Long-acting NPY analogues activating Y1R, Y4R, and Y5R receptors protect islets from cytotoxic damage [17]. Notably, selective Y4 receptor activation alone recapitulates these protective effects in vitro, indicating individual receptor subtypes may independently promote islet resilience [14]. Conversely, Y1R inhibition increases β-cell proliferation, whilst elevated NPY expression marks stress-exposed, dedifferentiated β-cells [18, 19]. These findings point to receptor-specific roles in regulating β-cell fate and adaptation, yet the cellular localization, molecular mechanisms, and therapeutic potential of individual NPY receptors remain unclear.

Among NPY receptor subtypes, Y4R is particularly compelling for therapeutic development. Selectively activated by PP and expressed in pancreatic islets, *Y4R* activation alone protects islets from inflammatory stress in vitro [12, 20]. Unlike *Y1R*, which broadly suppresses insulin secretion, Y4R may offer a more targeted strategy for enhancing β-cell resilience without disrupting glucose homeostasis. However, critical gaps persist: the precise cellular expression pattern of *Npy4r* within islets is unresolved, the transcriptional programmes and signalling networks mediating *Npy4r* effects are unknown; and whether *Y4R* activation modulates the islet immune microenvironment or alters disease progression in vivo remains untested. Mechanistic and translational progress has also been hampered by the lack of selective, pharmacologically optimized *Y4R* agonists suitable for in vivo studies.

Here, we address these gaps using K22, a long-acting, selective Y4R agonist with in vivo suitability[21]. Using a combination of cell sorting with qPCR, RNAscope, ligand-competition binding, functional islet assays, transcriptomics, and an adoptive transfer model of autoimmune diabetes, we investigated whether Y4R agonism can enhance β-cell resilience and alter islet-directed immunity, providing first-in-class in vivo evidence for Y4R as a candidate target for disease-modifying, β-cell-centred intervention in T1DResults

### Y4R is enriched on β-cells in mouse and human islets

To determine the cellular localisation of Y4R within pancreatic islets, we first confirmed *Npy4r* expression in whole human and mouse islets by quantitative PCR (Figure 1A, B respectively). In mouse islets, flow cytometric sorting of CD45-positive (immune) versus CD45-negative (endocrine) cell populations revealed *Npy4r* expression was markedly enriched in the CD45-negative fraction, consistent with predominant expression in islet endocrine cells (Figure 1C, D). To identify the specific endocrine cell type expressing *Npy4r*, we sorted β-cells from mouse islets using the zinc indicator DA-ZP1 (diacetylated Zinpyr1) [22], achieving a DA-ZP1-positive fraction of 87% with marked enrichment of insulin expression confirming high β-cell purity (Figure 1E, F, G). qPCR of sorted fractions demonstrated robust *Npy4r* expression in DA-ZP1-positive β-cells, whilst transcripts were undetectable in the DA-ZP1-negative fraction, directly establishing β-cells as the exclusively *Npy4r*-expressing endocrine cell type within mouse islets by this approach (Figure 1H). RNA *in situ* hybridization (RNAscope) confirmed *Npy4r* mRNA localization to individual β-cells within mouse islets (Figure 1I, J), consistent with low-moderate expression levels.

**Figure 1.**
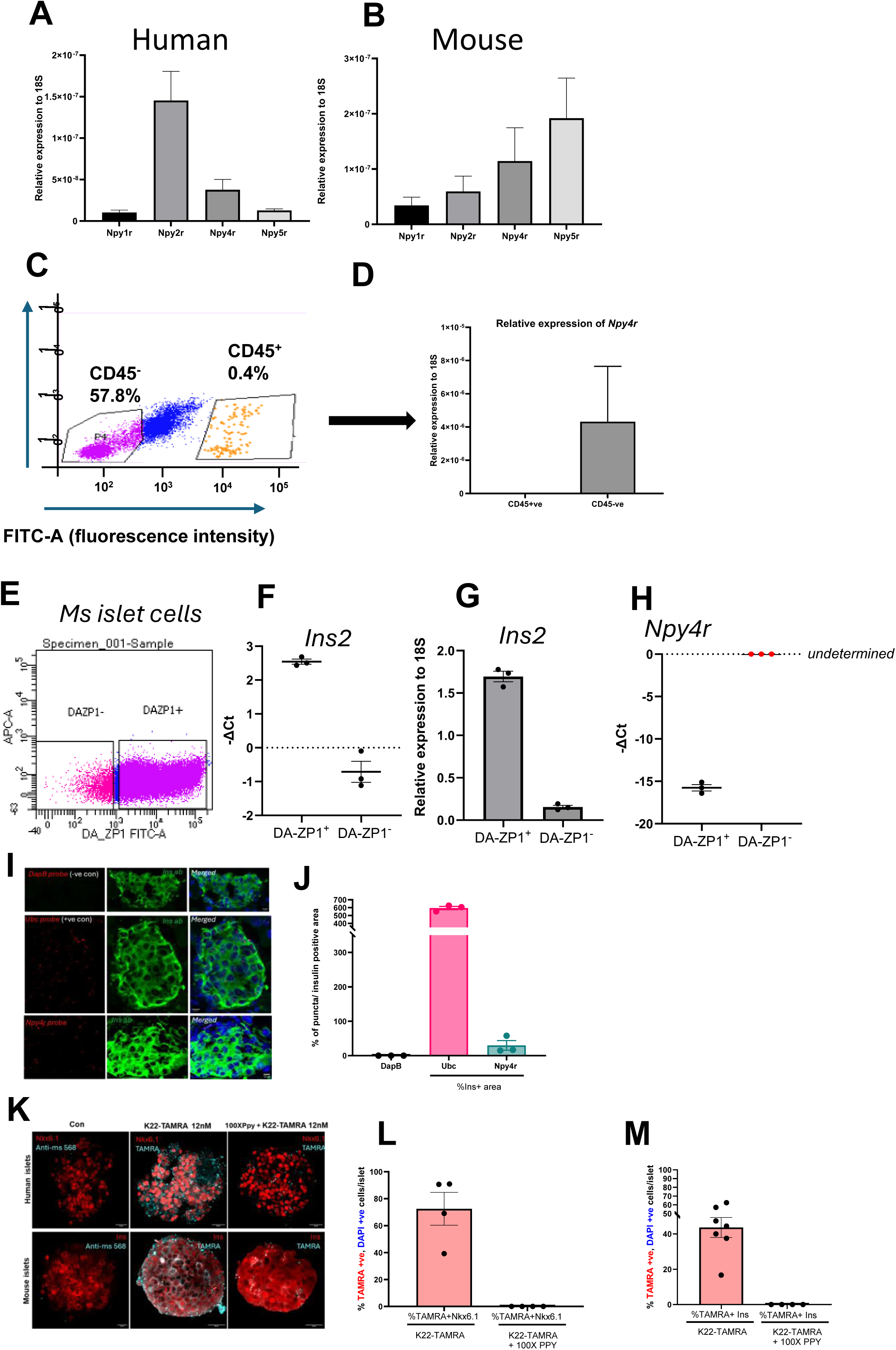
| Y4R is enriched in β-cells in mouse and human islets and is accessible to the selective agonist K22. (A, B) Relative mRNA expression of NPY1R/Npy1r, NPY2R/Npy2r, NPY4R/Npy4r and NPY5R/Npy5r in human (A) and mouse (B) islets, normalised to 18S rRNA (human, n = 100 islets per donor; mouse, n = 100 islets pooled from 6 mice; mean ± s.e.m.). (C) Representative flow-cytometry plot showing separation of CD45⁺ immune cells and CD45⁻ islet cells from mouse islets (n ≥ 1,800 islets pooled from 6 mice). (D) Relative expression of Npy4r (2⁻ΔΔCt, normalised to 18S) in whole islets, CD45⁺ and CD45⁻ fractions from mouse islets (n ≥ 1,800 islets pooled from 6 mice; mean ± s.e.m.), demonstrating enrichment of Npy4r in CD45⁻ cells. (E) DA-ZP1 staining of dispersed mouse islet cells used to enrich zinc-rich β-cells; the DA-ZP1⁺ gate contains 87.1% of events (n ≥ 1,800 islets from 6 mice). (F,G) qPCR validation of β-cell enrichment in DA-ZP1-sorted fractions showing Ins2 −ΔCt (F) and 2⁻ΔΔCt (G) in DA-ZP1⁺ versus DA-ZP1⁻ cells (n ≥ 1,800 islets from 6 mice); mean ± s.e.m.), with marked enrichment of Ins2 in DA-ZP1⁺ cells (mean ΔCt 2.54 vs −0.71). (H) Npy4r −ΔCt values in DA-ZP1⁺ and DA-ZP1⁻ fractions (n ≥ 1,800 islets from 6 mice; mean ± s.e.m.); Npy4r is readily detected in DA-ZP1⁺ cells (mean ΔCt 15.76) but is undetermined in DA-ZP1⁻ cells, indicating β-cell-biased expression. (I) RNAscope in situ hybridisation on mouse pancreatic sections showing negative-control DapB probe, positive-control Ubc probe and Npy4r probe (red), combined with insulin immunostaining (green) and DAPI (blue); Npy4r puncta localise to insulin-positive β-cells (n = 3 mice, ≥3 sections per mouse). (J) Quantification of RNAscope signal expressed as percentage of probe puncta per insulin-positive area for DapB, Ubc and Npy4r (n = 3 mice, 3 technical fields per mouse; mean ± s.e.m.). (K) Confocal images of human (top row) and mouse (bottom row) islets incubated with vehicle (Con), 12 nM K22–TAMRA, or 12 nM K22–TAMRA plus 100-fold excess pancreatic polypeptide (PP); K22–TAMRA fluorescence (cyan) co-localises with Nkx6.1 (human) or insulin (mouse) immunostaining (red) and is abolished by PP competition (human, n = 1 donors; mouse, n = 4 mice; ≥5 islets analysed per sample). (L,M) Quantification of K22–TAMRA binding in human (L) and mouse (M) islets, shown as the percentage of K22–TAMRA⁺, DAPI⁺ cells that are Nkx6.1⁺ (L) or insulin⁺ (M) per islet (human, n = X donors with ≥10 islets per donor; mouse, n = 5 mice with ≥10 islets per mouse; mean ± s.e.m.), with and without PP competition. K22–TAMRA robustly labels β-cells in both species and is competitively displaced by PP, supporting selective engagement of β-cell Y4R. Scale bars, 10 μm.

To confirm that *Y4R* protein is functionally accessible at the β-cell surface, we employed TAMRA-conjugated K22 (TAMRA-K22), a fluorescent derivative of our selective *Npy4r* agonist. As K22 binds *Npy4r* with high selectivity, TAMRA-K22 labelling of tissue sections provides a direct, antibody-independent readout of accessible Y4R at the cell surface. Incubation of mouse and human pancreatic sections with TAMRA-K22 revealed specific labelling of islet β-cells, confirmed by co-localisation with insulin and/or Nkx6.1 immunostaining (Figure 1K). To verify that this signal reflects *Y4R* -mediated binding rather than non-specific fluorescent labelling, sections were pre-incubated with excess pancreatic polypeptide (PP), the endogenous Y4R ligand, prior to TAMRA-K22 application. PP pre-incubation substantially blocked TAMRA-K22 signal, confirming receptor-mediated occupancy at the β-cell surface (Figure 1 K, L, M). Collectively, these findings establish that NPY4R is expressed in pancreatic β-cells in both mice and humans and is functionally accessible to selective *Y4R* agonists.

### *Npy4r* activation protects islets without impairing function

Having established *Npy4r* localisation to β-cells, we next evaluated the functional consequences of *Npy4r* activation using K22. We first assessed the safety profile of K22 across concentrations ranging from 6 to 500 nM in mouse islet cultures. K22 did not induce cytotoxicity at any concentration tested; notably, at 6–60 nM, K22 significantly reduced basal caspase-3/7 activity compared to vehicle control, indicating protection against culture-associated stress even in the absence of exogenous diabetogenic insults (Figure 2A). K22 did not affect glucose-stimulated insulin secretion in static assays across the full concentration range tested (60 pM–60 nM), with robust high-glucose responses maintained at levels comparable to untreated controls (Figure 2B). Dynamic perifusion confirmed that K22 (12 nM) did not alter insulin secretory kinetics in response to sequential low and high glucose challenges (Figure C-E). Single-islet calcium imaging demonstrated intact glucose-induced Ca²⁺ influx and ATP responsiveness, confirming preserved stimulus-secretion coupling (Figure 2F). K22 (12 nM) preserved both network connectivity and beta cell hub architecture, consistent with maintained coordinated islet activity (Figure 2G, H). Finally, glucose tolerance in K22-treated mice was unchanged compared with vehicle controls (Figure 2I), establishing that Y4R activation is neutral for whole-body glucose homeostasis at pharmacologically relevant concentrations.

**Figure 2.**
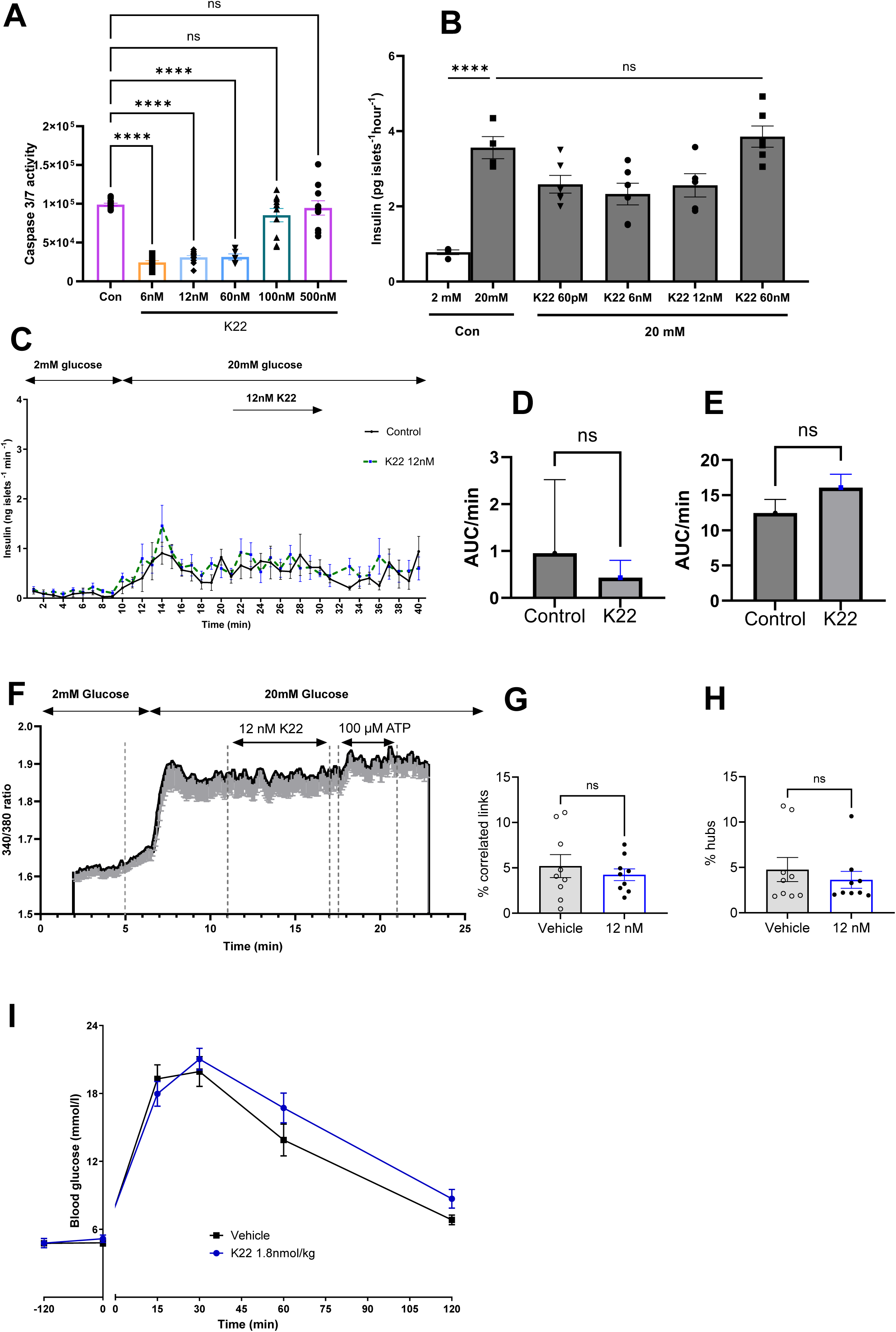
| K22 is non-toxic to β cells and does not impair insulin secretion *in vitro* or glucose tolerance *in vivo*. (A) Caspase 3/7 activity in human islets treated for 24 h with increasing concentrations of K22, expressed as relative light units; bars show mean ± s.e.m. with individual human islet preparations overlaid (n = 5-6 islets from 6 mice per group), and statistics indicate comparisons to vehicle control by one-way ANOVA with Dunnett’s multiple-comparison test. (B) Static insulin secretion from human islets stimulated with 2 mM or 20 mM glucose in the absence or presence of K22 (6–60 nM) for 24 h; bars represent mean insulin release per islet per hour ± s.e.m. (n = 4 human islets per group), with statistical comparisons performed by one-way ANOVA with Dunnett’s post hoc test versus 20 mM glucose alone. (C) Perifusion assay of human islets stimulated with 2 mM followed by 20 mM glucose with or without 12 nM K22, showing dynamic insulin secretion over time; traces represent mean ± s.e.m. of replicate perifusions (n = 4 islets per condition). (D,E) Quantification of first-phase (F, 0–12 min) and second-phase (G, 13–40 min) insulin secretion from the perifusion experiments in (E), expressed as AUC per minute; bars show mean ± s.e.m. (n = 4 islets per condition) and ns indicates no significant difference between control and K22 by paired two-tailed t-test. (F) Representative Fura-2 ratio trace from mouse islets stimulated with 2 mM then 20 mM glucose in the continued presence or absence of 12 nM K22, followed by 100 μM ATP as a positive control, illustrating preserved glucose-evoked Ca²⁺ oscillations (representative of n = 18-19 islets per preparation per condition). (G,H) Quantification of β cell functional connectivity from Ca²⁺ imaging in intact mouse islets exposed to vehicle or 12 nM K22, showing percentage of correlated links (B) and hub cells (C); data are mean ± s.e.m. with individual islets shown as dots (n = 9 islets from ≥3 mice), and ns indicates no significant difference by linear mixed-effects model. (I) Intraperitoneal glucose tolerance test in mice injected with vehicle or K22 (1.8 nmol per kg) 15 min before glucose challenge; blood glucose concentrations were measured over 120 min and are shown as mean ± s.e.m. (vehicle, n = 11; K22, n = 11). Glucose curves were analysed by two-way repeated-measures ANOVA with Sidak’s multiple-comparison test, revealing a significant effect of time but no main effect of treatment or time × treatment interaction, and no significant differences between vehicle and K22 at any time point, indicating that K22 does not alter glucose tolerance.

We next examined whether K22 protects islets against diabetogenic stressors. At concentrations as low as 6 nM, K22 pre-treatment substantially attenuated apoptosis induced by pro-inflammatory cytokines (IL-1β, TNF-α, IFN-γ) in both mouse and human islets, with dose-dependent reductions in caspase-3/7 activity across the 6–60 nM dose range (Figure 3A). This cytoprotective effect extended across mechanistically distinct diabetogenic insults, with K22 significantly reducing apoptosis induced by streptozotocin, palmitate, and thapsigargin in mouse islets (Figure 3B–D), suggesting engagement of a fundamental cellular resilience programme rather than blockade of a single death pathway. Immunofluorescence confirmed that K22 markedly reduced the frequency of caspase-3-positive cells in cytokine-exposed islets whilst preserving insulin-positive β-cell populations and islet morphology (Figure 3F, G). The translational relevance of these findings was underscored by equivalent protection in human islets, where K22 similarly attenuated cytokine-induced apoptosis (Figure 3E), demonstrating conservation of *Npy4r* -mediated cytoprotection across species.

**Figure 3.**
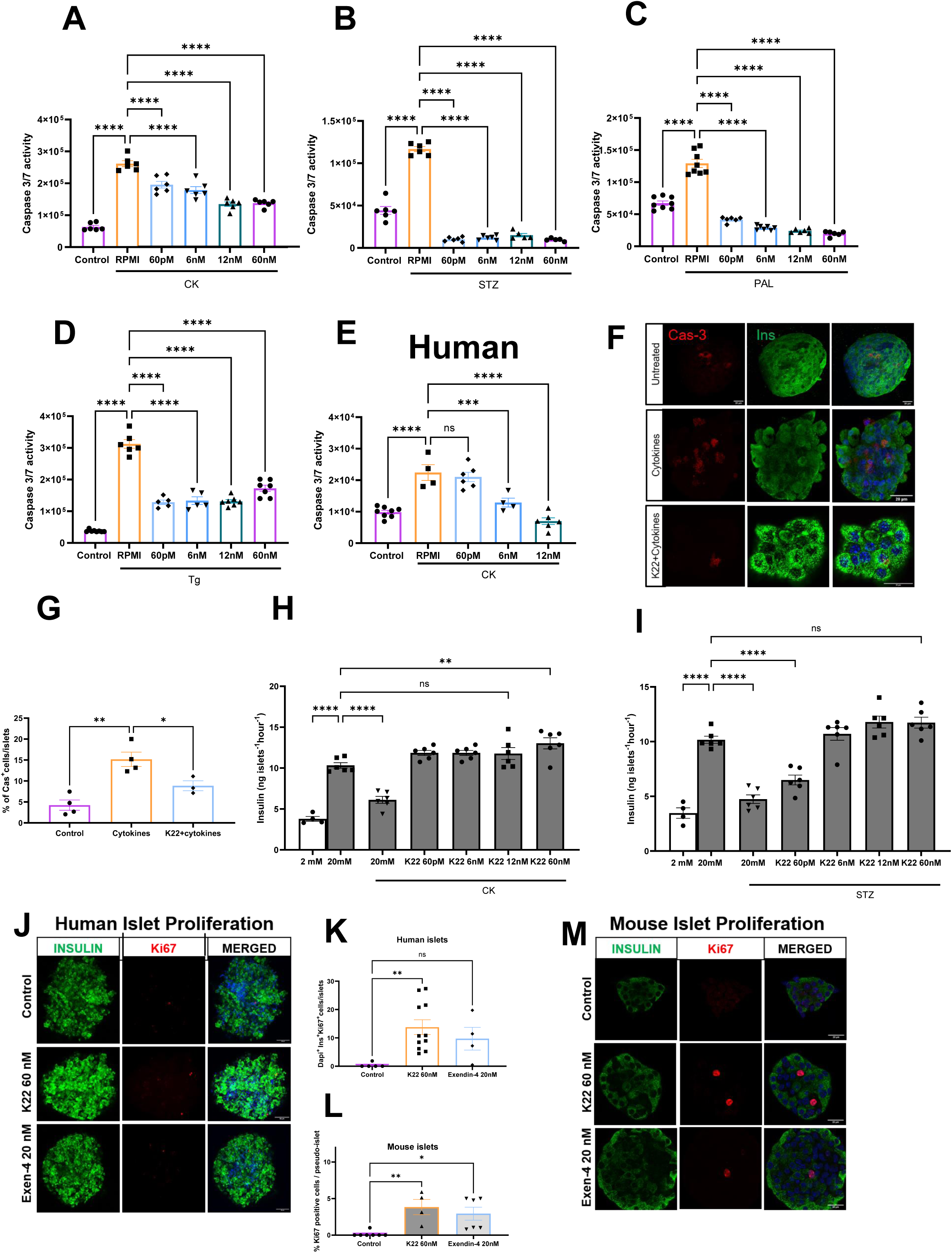
| K22 protects rodent and human β cells from apoptosis without impairing function and modestly enhances proliferation. (A–D) Caspase 3/7 activity in mouse islets cultured for 24 h in control medium, RPMI (vehicle for cytokines, CK), or RPMI containing CK, streptozotocin (STZ), palmitate (PAL) or thapsigargin (Tg) with or without K22 (6–60 nM), as indicated; bars show mean ± s.e.m. with individual preparations overlaid (n = 6-8 islets per group), and statistics indicate comparisons to the corresponding CK, STZ, PAL or Tg group by one-way ANOVA with Dunnett’s multiple-comparison test. (E) Caspase 3/7 activity in human islets exposed to CK with or without K22 (6–12 nM) for 24 h; data are mean ± s.e.m. in 1 donor (n = 4-6 islets per group), analysed by one-way ANOVA with Dunnett’s post hoc test versus CK alone. (F) Representative confocal images of mouse islets left untreated, treated with CK alone or CK plus 12 nM K22 and stained for cleaved caspase-3 (Cas-3, red), insulin (green) and DAPI (blue), illustrating reduced Cas-3⁺ β cells with K22; scale bars, 20 μm. (G) Quantification of Cas-3⁺ β cells per islet under the conditions in (F); each dot represents one islet (n = 3-4 islets from mixed pool of ≥3 mice) and bars indicate mean ± s.e.m., analysed by one-way ANOVA with Tukey’s multiple-comparison test. (H,I) Static insulin secretion from mouse islets pre-treated with CK (H) or STZ (I) for 24 h with or without K22 (6–60 nM) and then stimulated for 1 h with 2 mM or 20 mM glucose, as indicated; data are mean insulin release per islet per hour ± s.e.m. (n = 4-6 islets per group), analysed by one-way ANOVA with Dunnett’s test versus the corresponding stressed 20 mM glucose group. (J–M) Human (J,K) and mouse (L,M) islets cultured for 72 h in control medium, 60 nM K22 or 20 nM exendin-4 (Exen-4) and stained for insulin (green), Ki67 (red) and DAPI (blue); representative images are shown in (J) and (L), scale bars 20 μm. (K,M) Quantification of β cell proliferation expressed as percentage of DAPI⁺Ins⁺Ki67⁺ cells per islet (human, K) or percentage of Ki67⁺ cells per islet (mouse, M); bars represent mean ± s.e.m. with individual islets overlaid (human, n = 1 donor, 5-10 islets per group; mouse, n = 4-8 islets per group mixed pool of ≥3 mice), with significance between groups determined by one-way ANOVA with Dunnett’s multiple-comparison test.

Beyond preventing apoptosis, K22 preserved β-cell function under inflammatory and cytotoxic stress. Mouse islets exposed to pro-inflammatory cytokines or streptozotocin exhibited profound impairment of glucose-stimulated insulin secretion; K22 pre-treatment significantly restored glucose stimulated insulin release to levels comparable to unstressed controls (Figure 3H, I), demonstrating that *Npy4r* activation maintains functional β-cell capacity alongside structural integrity.

To investigate whether K22 influences β-cell mass, we assessed proliferation in both mouse and human islets. K22 (60 nM) significantly increased the frequency of Ki67-positive insulin-expressing cells in both species compared to vehicle controls (Figure 3J–M). In human islets, the increase in proliferating β-cells (DAPI⁺Ins⁺Ki67⁺) achieved levels comparable to the established proliferative agent Exendin-4 (20 nM), with equivalent findings in mouse. The coordinated enhancement of β-cell survival, function, and proliferation suggested that K22 engages transcriptional programmes governing β-cell identity and resilience, prompting comprehensive transcriptomic profiling to elucidate the underlying molecular mechanisms.

### Y4R agonism induces a β-cell resilience transcriptional programme

To identify the intrinsic transcriptional programmes mediating K22’s cytoprotective effects, we performed RNA sequencing on mouse islets treated with K22 or vehicle for 48 hours under basal culture conditions, in the absence of exogenous diabetogenic stress. This design reveals the molecular basis through which Y4R activation maintains β-cell health and resilience under the culture-associated stress inherent to isolated islet preparations.

Principal component analysis demonstrated clear separation between control and treated samples (PC1: 23.5% variance, PC2: 22.3% variance), indicating substantial transcriptional remodelling (Figure 4A); the residual variance not captured by PC1 and PC2 likely reflects inter-preparation heterogeneity inherent to primary islet biology. Differential expression analysis identified 360 genes at nominal p < 0.05 (217 upregulated, 143 downregulated), of which 63 remained significant after multiple testing correction (Benjamini–Hochberg adjusted p < 0.1). The nominal gene set was retained for downstream pathway analysis to maximise discovery sensitivity, with interpretation prioritising directional concordance across biologically coherent gene sets rather than individual gene significance, consistent with the IPA analytical framework applied throughout. Upregulated genes included core β-cell identity markers and secretory machinery (*Ins2*, *Chga*, *Vamp2*), the dense-core vesicle trafficking motor *Kif1a*, the chromatin modifier *Trim28*, and the antioxidant stress adaptation factor *Txnrd1*, whilst the most significantly downregulated gene was Reg3b, a stress-inducible antimicrobial peptide (Figure 4B). Marker-based enrichment analysis confirmed selective enhancement of β-cell identity gene sets (*Ins1*, *Ins2*, *Chga*, *Iapp*, *Nkx6-1*, *Pdx1*, *Ucn3*); whilst not all members of this gene set individually reached FDR correction thresholds, the coordinated directional enrichment across the β-cell signature was statistically significant, with minimal enrichment observed in gene sets marking other islet cell types, including α-cells (Gcg) and δ-cells (Sst) (Figure 4C). Canonical pathway analysis revealed coordinated activation of β-cell functional programmes including Insulin Processing (z-score = 2.0), Insulin Secretion Signalling (z-score = 1.73), and Synaptogenesis Signalling (z-score = 2.53), a pathway that shares core molecular components with dense-core vesicle fusion and trafficking machinery in β-cells, including *Kif1a*, *Vamp2*, *Rab3a*, *Syt7*, and *Syt13*. The KEAP1-NFE2L2 antioxidant pathway (z-score = 2.12) and Protein Ubiquitination pathway (z-score = 1.73) were co-ordinately activated, supporting enhanced cellular defence and proteostasis. In contrast, pathways linked to biosynthetic overload, EIF2 Signalling, Eukaryotic Translation Initiation, and rRNA Processing, were markedly suppressed, consistent with reduced biosynthetic and ER load in K22-treated islets. Modest enrichment of DNA synthesis and cell-cycle checkpoint pathways likely reflects adaptive DNA-damage and replication-stress responses rather than broad proliferative activation (Figure 4D).

**Figure 4.**
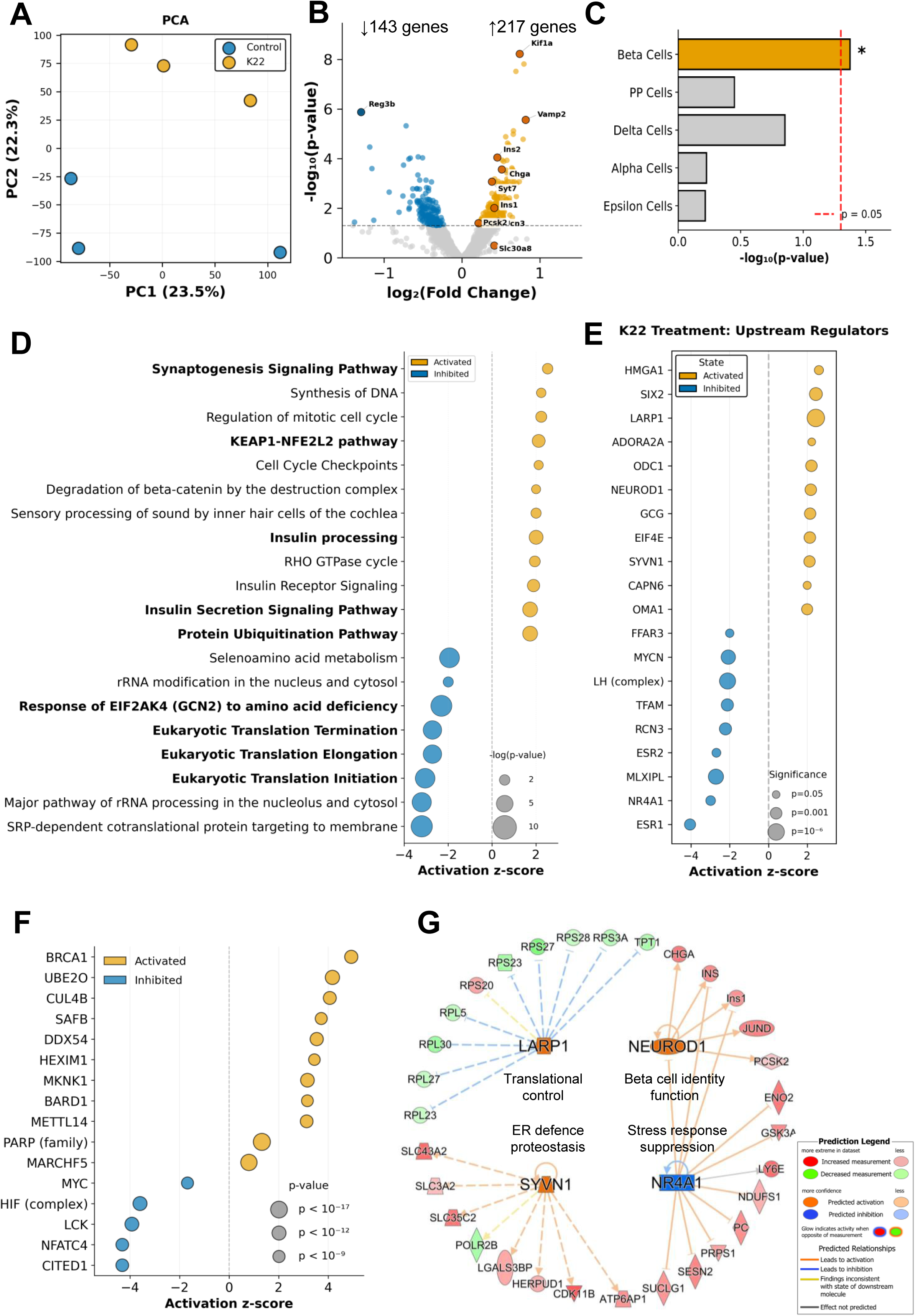
| K22 induces a β cell transcriptional programme that enhances identity, insulin processing and stress resistance. (A) Principal component analysis (PCA) of bulk RNA-seq from mouse islets treated for 24 h with vehicle (control) or K22, showing separation of conditions along PC1 and PC2 (1,500–1,800 islets per condition from a mixed pool of ≥6 mice). (B) Volcano plot of genes differentially expressed in K22-treated versus control islets, highlighting significantly upregulated (orange) and downregulated (blue) transcripts; selected β cell and stress-related genes are annotated. (C) Enrichment of endocrine cell-type signatures among genes upregulated by K22, showing over-representation of β cell identity genes (orange) compared with other islet cell types (grey); dashed red line indicates the significance threshold. (D) Canonical pathways significantly regulated by K22, plotted by activation z-score; orange circles denote activated and blue circles inhibited pathways, with circle size reflecting –log₁₀(p-value). (E) Predicted upstream regulators modulated by K22 treatment, ranked by activation z-score and significance; orange and blue indicate activated and inhibited regulators, respectively. (F) Additional transcriptional regulators affected by K22, including factors linked to DNA repair, RNA processing and hypoxia, displayed as in (E). (G) Schematic interaction network highlighting key K22-responsive upstream regulators (LARP1, NEUROD1, NR4A1, SYNV1) and their downstream targets involved in translational control, ER proteostasis, β cell identity and stress-response suppression; node colours indicate direction of regulation and predicted activation state.

Upstream regulator analysis predicted activation of key β-cell transcription factors including NEUROD1 (z-score = 2.19), SIX2 (z-score = 2.45), and HMGA1 (z-score = 2.61), consistent with enhanced β-cell maturation and preservation of the differentiated phenotype. Translational control mechanisms were predicted to be activated through LARP1 (z-score = 2.45) and EIF4E (z-score = 2.14). This pattern, activated translation regulators alongside suppressed global translation pathways, is consistent with a shift from bulk protein synthesis toward selective translation of stress-adaptive transcripts, a mechanism that would reduce ER burden whilst maintaining synthesis of essential survival proteins, though direct translational profiling will be required to confirm this interpretation. Proteostatic defences were predicted to be enhanced through SYVN1 (z-score = 2.12), an ER-associated E3 ubiquitin ligase, and OMA1 (z-score = 2.0), a mitochondrial stress protease. ADORA2A (z-score = 2.24), an anti-inflammatory G-protein coupled receptor, was also predicted to be activated, suggesting engagement of endogenous anti-inflammatory signalling.

Several transcriptional regulators showed predicted reduction in activity consistent with an overall reduction in cellular stress burden. NR4A1 (z-score = −2.99), an immediate-early stress-responsive transcription factor, showed suppressed predicted activity, likely reflecting the lower basal stress state of K22-treated islets relative to vehicle-treated controls undergoing culture-induced stress. MLXIPL/ChREBP (z-score = −2.71) and MYCN (z-score = −2.07), regulators of lipogenic gene expression and ribosomal protein synthesis respectively, were similarly inhibited, consistent with reduced biosynthetic burden (Figure 4E).

Causal network analysis predicted activation of DNA repair and chromatin remodelling regulators (BRCA1, CUL4B, BARD1), the m⁶A RNA methylation writer METTL14 (z-score = 3.13), which has established roles in β-cell survival and pancreatic cell differentiation, and stress adaptation enzymes (UBE2O, MKNK1). Predicted inhibited master regulators included NFATC4, LCK, the HIF complex, and MYC, consistent with coordinated dampening of inflammatory signalling and stress-amplifying programmes (Figure 4F).

Collectively, Y4R activation drives coordinated transcriptional reprogramming characterised by enhanced β-cell identity programmes (NEUROD1, SIX2, HMGA1), activation of insulin biosynthesis and secretion pathways, antioxidant responses (KEAP1-NFE2L2), proteostatic defences (SYVN1, ubiquitination), and mitochondrial quality control (OMA1), alongside suppression of global translational burden and stress-responsive transcription factors (NR4A1, MLXIPL, MYCN) (Figure 4G). This coordinated transcriptional signature is consistent with the observed reduction in caspase-3/7 activity and improved islet survival under culture stress, and provides a molecular framework for the broad cytoprotection demonstrated across multiple diabetogenic stressors.

### NPY4R agonism suppresses chemokines and limits immune recruitment

Transcriptomic analysis revealed multiple lines of evidence suggesting Y4R activation may modulate the islet immune microenvironment: predicted activation of the anti-inflammatory receptor ADORA2A and inhibition of master regulators of immune activation and T cell signalling, NFATC4 and LCK, alongside broad attenuation of pro-inflammatory transcriptional signatures. Although this transcriptional profiling was performed under basal culture conditions, we next investigated whether equivalent immunomodulatory effects operate under cytokine challenge by directly measuring inflammatory mediator output from K22-treated islets exposed to pro-inflammatory stress.

We first assessed CXCL10, a critical chemokine that recruits autoreactive T cells to pancreatic islets and drives insulitis progression [23]. Quantification by ELISA revealed that islets exposed to pro-inflammatory cytokines (IL-1β, TNF-α, IFN-γ) exhibited robust CXCL10 induction; K22 dose-dependently suppressed this response, nearly abolishing cytokine-induced upregulation at concentrations as low as 6 nM, with treated islets exhibiting CXCL10 levels comparable to unstimulated controls (Figure 5A). To determine the breadth of K22’s immunomodulatory effects, we employed multiplex immunoassay analysis of 26 analytes. K22 blunted cytokine-driven production of inflammatory mediators with established roles in diabetes pathogenesis: CXCL10 orchestrates recruitment of autoreactive CD8⁺ T cells that directly destroy β-cells; CCL3, CCL4, and CCL7 coordinate monocyte and macrophage infiltration into islets; CXCL2 recruits neutrophils that mediate early inflammatory responses; IL-6 sustains the pro-inflammatory milieu and directly impairs β-cell function; and IL-5 promotes eosinophil-mediated inflammation (Figure 5B–H). In contrast, CCL2 was modestly increased (25%, p < 0.05), the biological significance of which is currently unclear, whilst IL-2, essential for regulatory T cell maintenance, was also elevated (Figure 5I, J). Other analytes in the panel (IL-9, IL-10, IL-12p70, IL-17A, IL-18, IL-22, IL-23, IL-27, CCL5, GM-CSF, CXCL1) were either not induced by cytokine treatment or unaffected by K22 co-treatment (Figure S1). Collectively, K22 remodels the islet inflammatory environment, suppressing chemokines and cytokines that recruit and activate pathogenic immune cells whilst preserving signals that support immune regulation.

**Figure 5.**
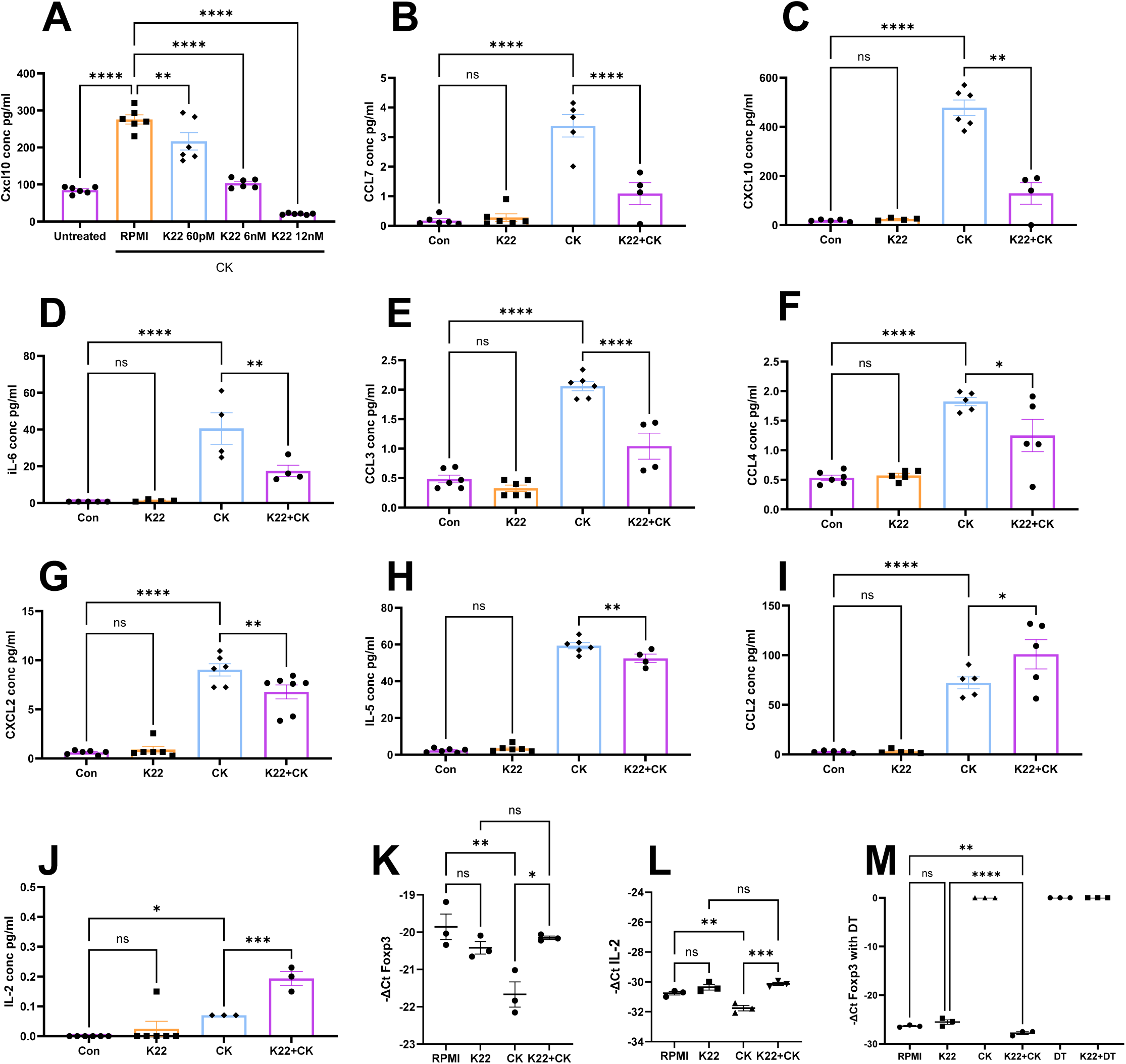
| K22 suppresses cytokine-induced inflammatory mediator production while preserving IL-2 and Foxp3-associated regulatory signals. (A) CXCL10 protein concentrations measured by ELISA from mouse islets cultured for 24 h under basal conditions (Untreated/RPMI) or with a pro-inflammatory cytokine cocktail (CK) in the absence or presence of K22 at the indicated concentrations; bars show mean ± s.e.m. with technical replicates (n=3-6) overlaid (1,500–1,800 islets per group from mixed pools of ≥6 mice). (B–H) CXCL10 (B), CCL3 (C), CCL4 (D), CCL7 (E), CXCL2 (F), IL-6 (G) and IL-5 (H) measured by multiplex bead assay in lysates and/or supernatants from mouse islets cultured for 24 h in control medium (Con/RPMI), K22 alone, CK alone or CK plus K22 (K22+CK), as indicated; bars represent mean ± s.e.m. with individual preparations shown as dots (same pooled-islet design as in A), and statistical comparisons between conditions are indicated (ns, not significant). (I) CCL2 concentrations in islet supernatants from the same culture conditions as in B–H, illustrating partial suppression of cytokine-induced CCL2 release by K22; data are mean ± s.e.m. with technical replicates overlaid. (J) IL-2 protein levels in supernatants from mouse islets cultured under the indicated conditions, demonstrating preservation or enhancement of IL-2 in the presence of K22 during cytokine exposure; bars show mean ± s.e.m. with individual replicates plotted. (K,L) Foxp3 (K) and Il2 (L) mRNA expression in islets cultured for 24 h in RPMI, K22 alone, CK alone or K22+CK, quantified by qPCR and expressed as −ΔCt relative to housekeeping genes; symbols denote individual cultures and horizontal lines indicate mean ± s.e.m. (M) Foxp3 expression (−ΔCt) in Foxp3-DTR islets treated under the same conditions as in K and L, with additional diphtheria toxin controls (DT and K22+DT), confirming DTR-dependent Foxp3 depletion and preservation of Foxp3 expression by K22 during cytokine exposure; data are mean ± s.e.m. with technical replicates overlaid, and significance between groups is indicated (ns, not significant).

Reduced IL-6 together with increased IL-2 in K22-treated islets suggested a shift toward a more regulatory immune environment, as IL-6 can destabilise FoxP3 expression and IL-2 supports maintenance of regulatory T cells (Figure 5D, J). We therefore assessed *Foxp3* and *Il-2* expression by qPCR. Pro-inflammatory cytokines markedly reduced *Foxp3* transcript levels in islets, whereas K22 co-treatment preserved Foxp3 expression near baseline levels (Figure 5K). *Il-2* expression was similarly maintained in K22-treated islets compared to vehicle controls (Figure 5L). Consistent with this, in Foxp3 reporter islets, K22 increased the proportion of FoxP3 reporter-positive cells during cytokine exposure (Figure 5M). Together, these data suggest preservation of FoxP3-expressing cells under inflammatory stress. The cellular identity and functional state of these FoxP3-positive cells, and the source of IL-2 in these preparations, remain to be determined. Accordingly, these findings support preservation of pro-regulatory markers under inflammatory stress but do not by themselves establish functional tolerance induction.

To validate these molecular signatures functionally, we assessed immune cell recruitment using reformed islet co-culture systems with haplotype-matched immune cells. Transwell migration assays demonstrated that K22 significantly attenuated cytokine-induced CD8⁺ T cell and CD4⁺ T cell migration toward islets (Figure 6A, B) and reduced both RAW264.7 macrophage and primary peritoneal macrophage chemotaxis (Figure 6C, D), consistent with the coordinated suppression of CXCL10 and CCL chemokines. Confocal imaging of reformed and native islets revealed that cytokine treatment resulted in extensive loss of insulin-positive β-cells, marked CD80⁺ macrophage infiltration, and architectural destruction in both mouse and human islet preparations. In contrast, K22-treated islets maintained robust insulin staining, exhibited reduced immune cell infiltration, and preserved structural integrity at both 2h and 15-20h timepoints (Figure 6F, D, F-L). These functional outcomes demonstrate that K22’s coordinated suppression of inflammatory mediators and preservation of *Foxp3* expression translate to reduced immune cell recruitment and protection of islet architecture under inflammatory assault.

**Figure 6.**
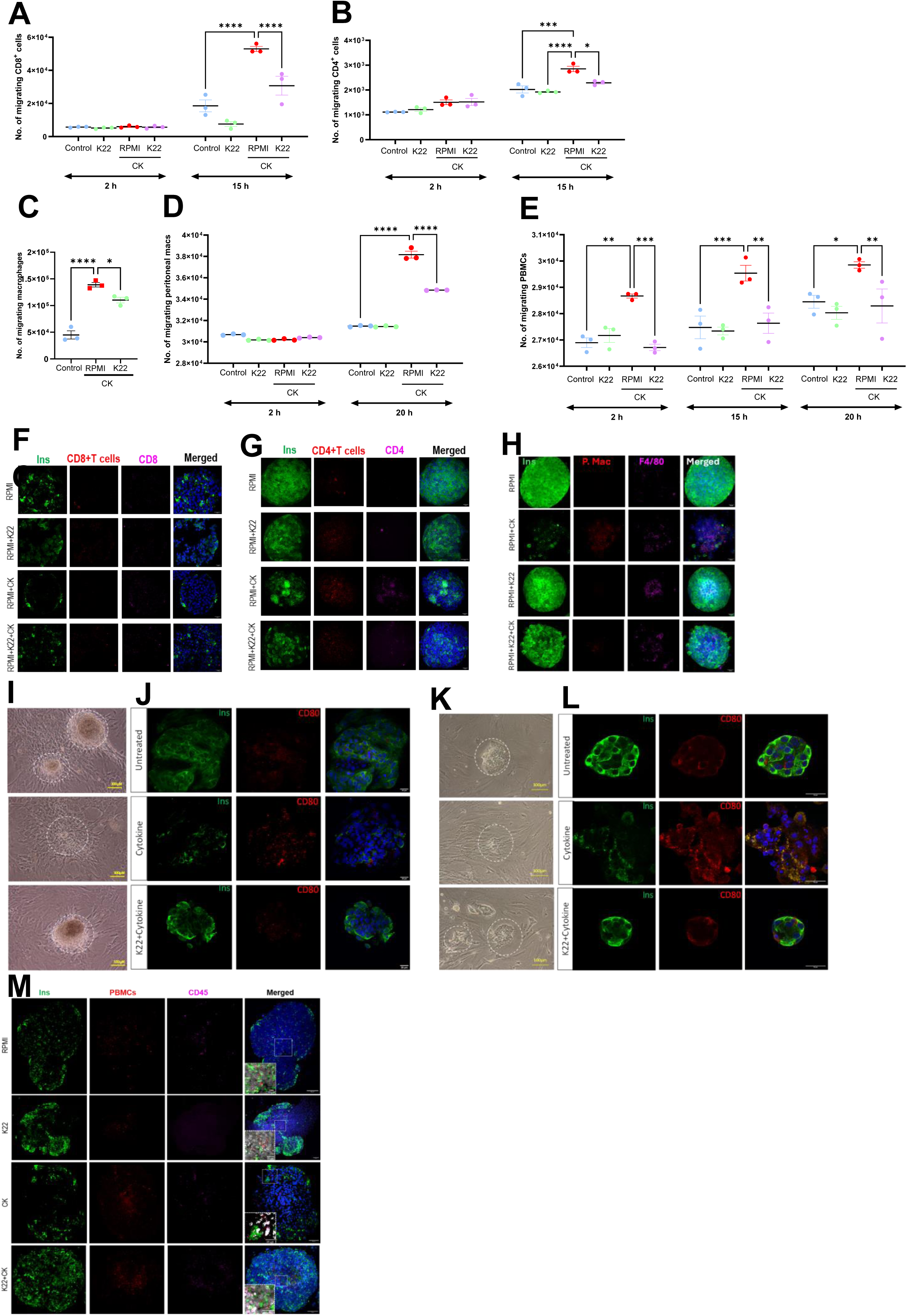
| K22 reduces cytokine-driven recruitment of T cells and macrophages to mouse and human islets and preserves β cell morphology and costimulatory signalling. (A,B) Quantification of migrating CD8⁺ (A) and CD4⁺ (B) T cells in Transwell assays in which T cells were exposed for 2 h or 15 h to conditioned media from mouse islets treated with control medium (2% FBS, 1% P/S), K22 alone, cytokines in RPMI (CK, RPMI) or cytokines plus K22 (K22+Cytokine); islets in each experiment were prepared from a mixed pool of 6 mice (100 islets per condition), with 3 technical Transwell replicates per condition (n = 3 wells per condition), and bars show mean ± s.e.m. numbers of migrated cells with individual wells overlaid and statistical comparisons indicated. (C) Quantification of migrating macrophages (RAW264.7 cells) in a Transwell assay in which macrophages were exposed for 20 h to conditioned media from mouse islets treated with control medium (Con), cytokines (CK) or cytokines plus K22 (K22+Cytokine); islets were from a mixed pool of 6 mice with 3 technical replicates per condition (n = 3 wells per condition), and bars represent mean ± s.e.m. of migrated macrophages with individual wells overlaid and statistical comparisons indicated. (D) Quantification of migrated cytokine-treated peritoneal macrophages in a Transwell invasion assay performed with mouse islets from a mixed pool of 6 mice; peritoneal macrophages were exposed for 2 h or 20 h to islets treated with control medium, cytokines, or cytokines plus K22 (K22+Cytokine) in 3 technical replicates per condition (n = 3 wells per condition), and bars show mean ± s.e.m. numbers of invading macrophages with individual wells overlaid and statistical comparisons indicated. (E) Quantification of migrated, activated human PBMCs in a Transwell invasion assay performed with human islets, in which PBMCs were exposed for 2 h, 15 h and 20 h to islets treated with control medium, cytokines, or cytokines plus K22 (K22+Cytokine); n = 1donor with 3 technical replicates per condition, and bars show mean ± s.e.m. numbers of invading PBMCs with individual wells overlaid and statistical comparisons indicated. (F,G) Representative confocal images of mouse islets co-cultured with CD8⁺ (F) or CD4⁺ (G) T cells under the indicated conditions (RPMI, RPMI+K22, RPMI+CK, RPMI+K22+CK); images show insulin (Ins, green), seeded CD8⁺ or CD4⁺ T cells in the upper Transwell insert (red), CD8 or CD4 cells within islets (magenta) and DAPI (blue) (n = 3 experiments, ≥5 islets imaged per condition), illustrating cytokine-induced accumulation of CD8⁺ and CD4⁺ T cells within islets and its attenuation by K22. (H) Representative confocal images from the 20 h peritoneal macrophage–mouse islet invasion assay (n = 3 experiments, ≥5 islets per condition), showing Qtracker-labelled peritoneal macrophages infiltrating cytokine-treated islets and reduced invasion in the presence of K22, with Ins (green), macrophages (F4/80 or Qtracker, magenta) and DAPI (blue). (I,J) Phase-contrast (I) and immunofluorescence (J) images of human islets cultured under basal conditions (Untreated), with cytokines alone, or cytokines plus K22 (K22+Cytokine); phase-contrast images show islet morphology (dashed circles), and confocal images (n = 3 donors, ≥5 islets per condition) show Ins (green), CD80 (red) and DAPI (blue), with cytokines disrupting insulin staining and increasing CD80, whereas K22 preserves islet morphology and reduces CD80 signal. (K, L) Corresponding phase-contrast (K) and immunofluorescence (L) images of mouse islets under the same conditions (Untreated, Cytokine, K22+Cytokine; n = 3 experiments, pooled islets from 6 mice, ≥5 islets per condition), showing similar protection of mouse islet morphology and suppression of cytokine-induced CD80 expression by K22. (M) Representative confocal images from the 15 h human PBMC–human islet invasion assay (n = 1 donor, ≥5 islets per condition), showing Qtracker-labelled PBMCs infiltrating cytokine-treated human islets and reduced invasion in the presence of K22, with Ins (green), PBMCs (magenta) and DAPI (blue) outlining islet structure.

To determine whether K22’s suppression of islet chemokine output translates to reduced immune cell recruitment in a fully human system, IL-2-activated, T cell-enriched human PBMCs were assessed for migration and invasion toward cytokine-stressed human islets in matched Transwell assays. Cytokine treatment of human islets significantly increased PBMC migration at all timepoints assessed (2 h, 15 h, and 20 h), and K22 co-treatment consistently and significantly attenuated this response, returning migration to levels comparable to unstimulated controls at each timepoint (Figure 6E). Complementary invasion assays performed at 2 h, 15 h, and 20 h showed progressive accumulation and penetration of activated PBMCs into cytokine-treated human islets, with markedly reduced peri-islet and intra-islet invasion in K22-treated cultures, mirroring the migration data (Figures 6M and S2A, B). As PBMCs were not HLA-matched to the islet donors, this model principally reports cytokine-driven rather than antigen-specific immune recruitment; nonetheless, these findings demonstrate that K22’s immunomodulatory effect is preserved in a fully human immune–islet setting.

Consistent with these findings, imaging of Qtracker-labelled PBMC imaging confirmed early peri-islet association at 2 h and extensive intra-islet invasion by 20 h under cytokine conditions, both markedly reduced by K22 co-treatment (Figure S2). Taken together, Y4R agonism couples intrinsic β-cell cytoprotection with suppression of islet chemokine production and consequent limitation of CD8⁺ T cell, CD4⁺ T cell, macrophage, and human PBMC recruitment in both mouse and fully human immune-islet systems, whilst preserving pro-regulatory markers within the islet microenvironment, prompting us to test whether K22 can delay diabetes progression in vivo.

### NPY4R agonism delays accelerated diabetes onset

To evaluate therapeutic efficacy in autoimmune diabetes, we employed an adoptive transfer model in which Rag1-deficient mice received in vitro islet autoantigen-activated NY8.3 CD8⁺ T cells, circumventing endogenous adaptive immune regulation to isolate the effects of K22 on disease progression (Figure 7A). Systemic K22 treatment significantly delayed diabetes onset compared to saline-treated controls (n = 8 K22-treated, n = 12 saline controls, log-rank test p = 0.0372, Figure 7B). Median time to diabetes was approximately 11 days in K22-treated mice versus 7 days in controls, with all mice in both groups reaching diabetic endpoint by day 14. Whilst K22 did not prevent diabetes in this aggressive model, the significant delay in onset demonstrates that Y4R agonism can meaningfully slow autoimmune disease progression even against a defined, highly diabetogenic CD8⁺ T cell challenge.

**Figure 7.**
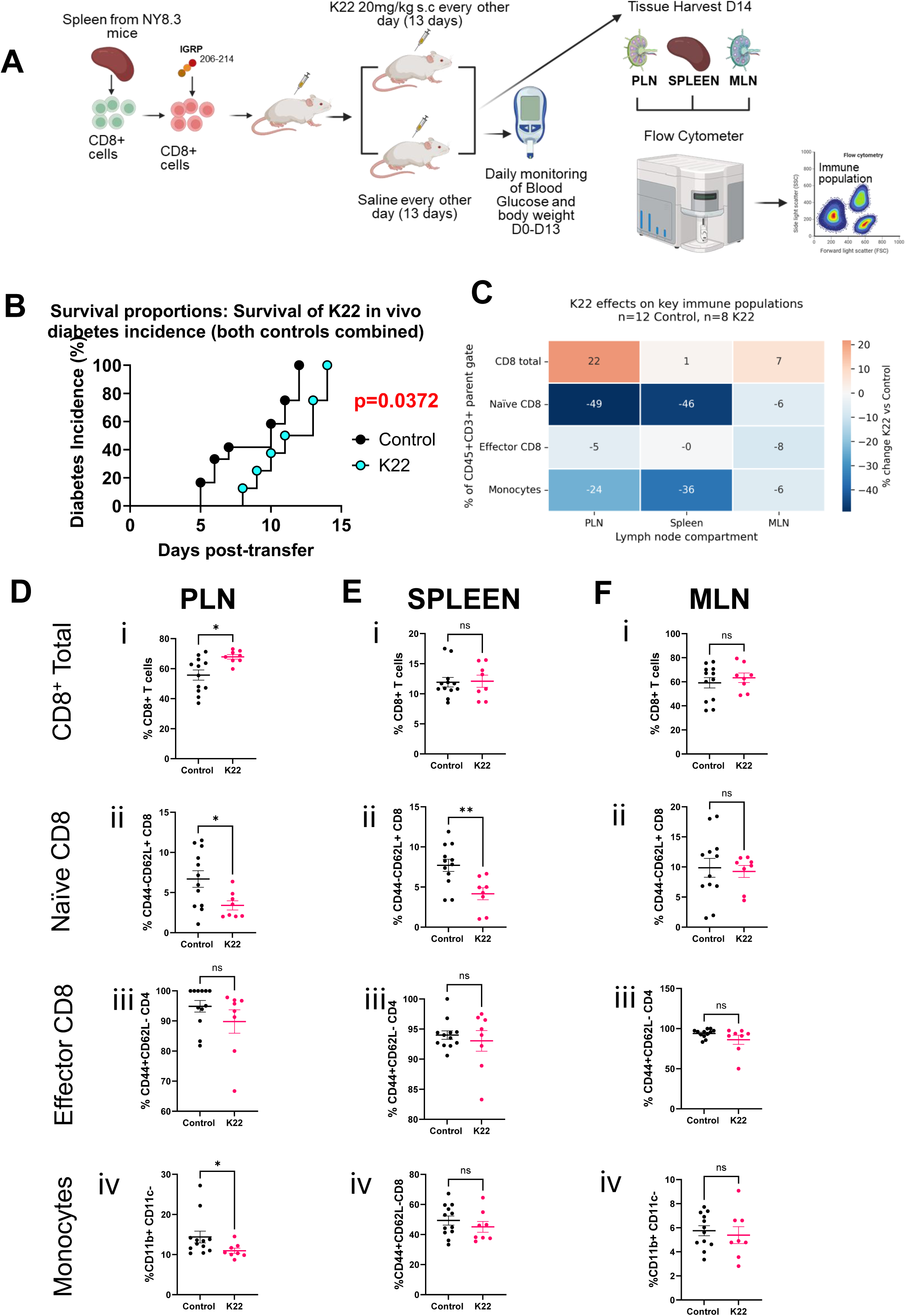
| K22 delays diabetes onset in an adoptive transfer model without causing broad systemic immune perturbation. (A) Schematic of the NY8.3→Rag1⁻/⁻ adoptive-transfer experiment: NY8.3 CD8⁺ T cells are activated with IGRP₂₀₆–₂₁₄ and transferred into Rag1-deficient NOD recipients, which then receive K22 (20 mg/kg s.c. every other day for 13 days) or saline control; blood glucose and body weight are monitored daily and peripheral lymph nodes (PLN), spleen and mesenteric lymph nodes (MLN) are harvested on day 14 for flow cytometry (n = 8 mice per group). (B) Kaplan–Meier plot of cumulative diabetes incidence in control- versus K22-treated mice, with diabetes defined as two consecutive blood glucose measurements above threshold; the p value was determined by log-rank test. (C) Heat map summarising percentage change in key immune populations in K22-treated versus control mice across PLN, spleen and MLN (total CD8⁺ T cells, naïve CD8⁺ T cells, effector CD8⁺ T cells and CD11b⁺CD11c⁻ monocytes). (D–F) Flow-cytometric quantification of total CD8⁺ T cells, naïve CD8⁺ T cells (CD44⁻CD62L⁺), effector CD8⁺ T cells (CD44⁺CD62L⁻) and CD11b⁺CD11c⁻ monocytes in PLN (D), spleen (E) and MLN (F) from control- and K22-treated mice at endpoint; each symbol represents an individual mouse, bars denote mean ± s.e.m., and statistical comparisons between groups are indicated (ns, not significant), showing reduced naïve CD8⁺ T cells and monocytes in PLN with K22 but no evidence of widespread immune dysregulation across secondary lymphoid organs.

To characterise immune changes associated with K22 treatment, we performed flow cytometric analysis of the spleen, pancreatic draining lymph nodes (PLN), and mesenteric lymph nodes (MLN) (Figure 7D). In the PLN, the primary site of islet antigen presentation and autoreactive T cell priming, K22-treated mice exhibited an increased frequency of CD8⁺ T cells together with a reduction in naïve (CD44⁻CD62L⁺) CD8⁺ T cells (Figure 7C, D i, ii). A similar reduction in CD44⁻CD62L⁻ CD8⁺ T cells was observed in the spleen (Figure 7C, E ii). A similar trend was observed in the MLN, though this did not reach statistical significance (p = 0.5714, Figure 7F ii). These data indicate altered CD8⁺ T cell phenotype and compartmental distribution within draining and systemic lymphoid tissue. Whether this reflects changes in priming, retention, or egress will require further investigation. While purity for the CD8⁺ T cell transfer exceeded 95%, low-level CD4⁺ T cells were detectable in recipient mice, however, CD4⁺ T cell subpopulations including central memory, effector subsets, total CD4⁺ frequencies, and regulatory T cell percentages remained unchanged across all compartments (Figure S3 A-C).

To assess whether K22 directly modulates autoreactive CD8⁺ T cell activation, we examined antigen-driven responses in isolated NY8.3 CD8⁺ T cells stimulated with their cognate antigen peptide IGRP₂₀₆₋₂₁₄ presented by splenic antigen-presenting cells. Npy4r transcripts were detectable and increased following antigen activation, indicating that Y4R expression may be inducible in activated CD8⁺ T cells (Figure 8A). However, across IGRP peptide concentrations, K22 did not alter CD8⁺ T cell proliferation and had no detectable effect on IFNγ or MIP1β secretion, with only a modest increase in TNFα (Figure 8B–F). Although Y4R is expressed by activated autoreactive T cells, K22 does not broadly suppress their effector programming under these conditions, supporting an islet-centred mechanism as the primary contributor to the in vivo phenotype rather than direct T cell suppression.

**Figure 8.**
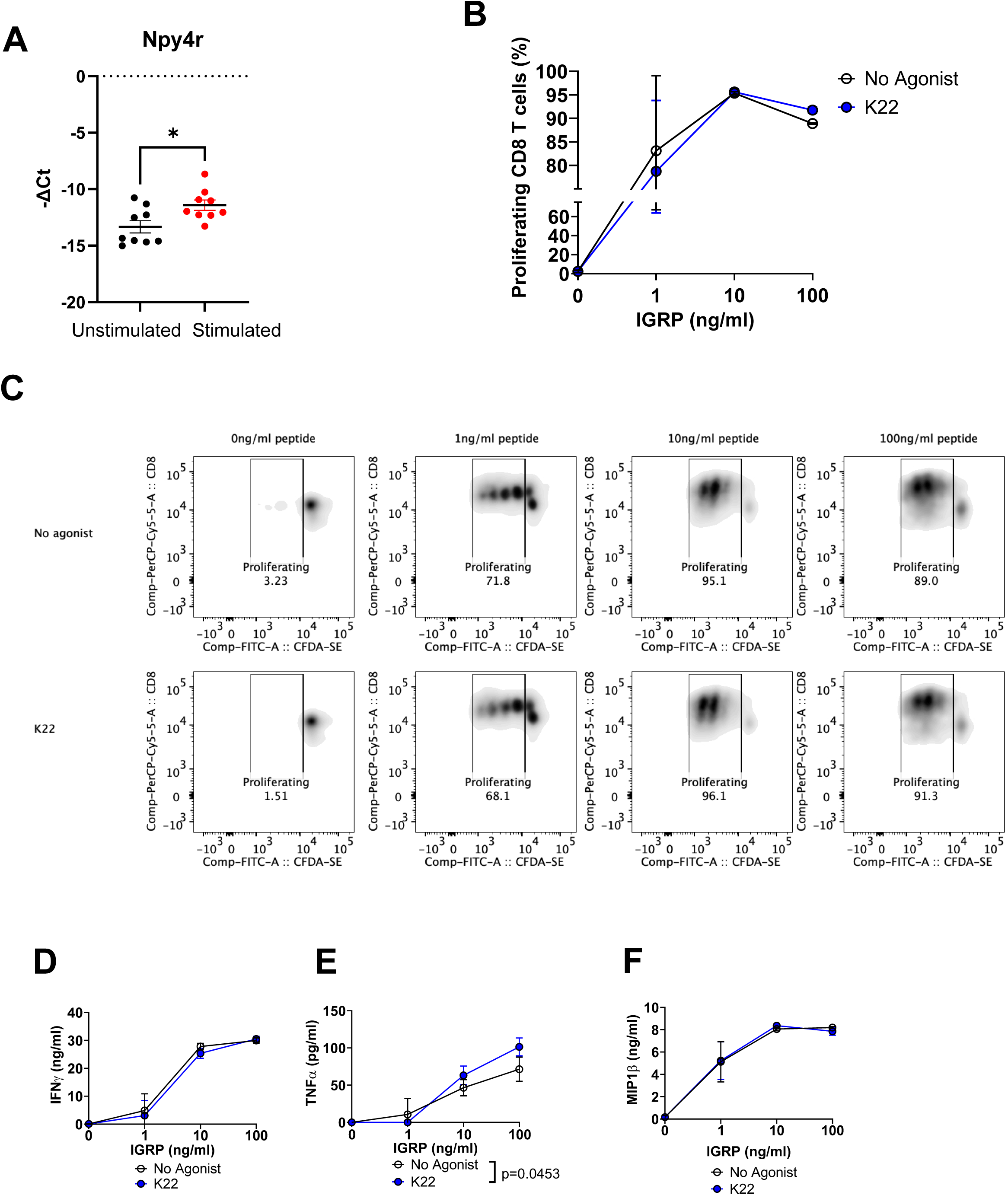
|: K22 does not impair Npy4r expression or antigen-specific activation of diabetogenic CD8⁺ T cells. (A) Npy4r mRNA expression in CD8⁺ T cells left unstimulated or stimulated through the T-cell receptor, quantified by qPCR and expressed as −ΔCt relative to housekeeping genes; each dot represents an independent sample from a mixed pool of >3 mice per group, and bars show mean ± s.e.m., with statistical significance indicated. (B) Proliferation of IGRP-specific CD8⁺ T cells cultured with increasing concentrations of IGRP peptide (0–100 ng ml⁻¹) in the presence of no agonist or K22, assessed by CFDA-SE dilution; data are mean ± s.e.m. percentage of proliferating cells from pooled T cells derived from >3 mice per group. (C) Representative flow-cytometry plots of CFDA-SE versus CD8 for IGRP-specific CD8⁺ T cells stimulated with the indicated IGRP peptide concentrations in the absence (top row) or presence (bottom row) of K22, with the percentage of proliferating cells annotated in each panel. (D–F) IFNγ (D), TNFα (E) and MIP1β (F) secretion by IGRP-specific CD8⁺ T cells stimulated with increasing IGRP peptide doses in the presence of no agonist or K22, measured in culture supernatants; values represent mean ± s.e.m. from T cells pooled from >3 mice per group, with statistically significant differences between conditions indicated.

Myeloid populations in the PLN were also altered in K22-treated mice, with decreased CD11b⁺CD11c⁻ cells and increased CD11b⁻CD11c⁺ dendritic cells (Figure 7D iv, S3 x), consistent with changes in myeloid composition within the draining node, though the functional significance of this shift requires further investigation.

Collectively, systemic K22 delayed accelerated autoimmune diabetes onset in this stringent NY8.3 CD8⁺ T cell adoptive-transfer model, providing the first in vivo evidence that Y4R agonism can be therapeutically targeted to delay autoimmune diabetes progression. The in vivo efficacy aligns with our ex vivo findings that K22 enhances β-cell stress resilience, reduces chemokine-driven immune cell recruitment toward cytokine-exposed islets, and preserves pro-regulatory markers within the islet microenvironment. Together, these dual protective actions, enhancing β-cell resilience whilst dampening local immune attack, establish Y4R as a promising therapeutic target for type 1 diabetes that operates through a mechanistically distinct approach to current immunosuppressive strategies.

## Discussion

Type 1 diabetes remains a chronic autoimmune disorder without a cure, and current disease-modifying strategies face important limitations[1, 24]. The approval of teplizumab as the first disease-modifying therapy shown to delay progression to clinical T1D represents a major advance [6]. However, as a systemically acting anti-CD3–mediated T cell–modulating therapy, its use is accompanied by immune-related risks that may constrain broad or repeated application, particularly in paediatric populations [5]. Other immunomodulatory approaches, including JAK inhibition, similarly depend on systemic immune modulation [25, 26]. Conversely, strategies focused solely on β-cell replacement or regeneration do not address ongoing autoimmune destruction of endogenous or transplanted β-cells [27, 28]. An ideal therapeutic approach would therefore (i) strengthen intrinsic β-cell resilience and (ii) locally attenuate islet inflammatory recruitment signals, limiting immune-mediated damage without requiring broad systemic immunosuppression [29]. Here, we identify Y4R agonism as a candidate strategy that engages both objectives: K22 confers direct β-cell cytoprotection through coordinated transcriptional reprogramming while simultaneously suppressing islet chemokine production and preserving local pro-regulatory features, collectively aligning with the observed delay in autoimmune diabetes onset in vivo.

Our data establish Y4R as a direct mediator of β-cell protection. Previous studies using antibody-based approaches have reported variable Y4R localisation within islets, a challenge compounded by the recognised limitations of GPCR antibody specificity [30]. Using complementary approaches including cell sorting coupled with qPCR, RNAscope, and receptor binding with TAMRA-conjugated K22, we demonstrate robust Y4R expression in insulin-positive β-cells in both mouse and human islets. The TAMRA-K22 signal was substantially blocked by pre-incubation with pancreatic polypeptide, providing antibody-independent functional evidence that Y4R is present and accessible on β-cells. This localisation establishes β-cells as the primary cellular target of Y4R agonists, providing a mechanistic basis for the cytoprotective effects observed with K22. The functional consequence of this β-cell expression was validated across multiple experimental systems: K22 protected against inflammatory cytokines, streptozotocin, palmitate, and thapsigargin whilst preserving glucose-stimulated insulin secretion and enhancing proliferation. These effects were conserved in human islets, underscoring translational relevance. The breadth of protection across mechanistically distinct stressors, inflammatory, cytotoxic, lipotoxic, and ER stress, suggests that Y4R activation engages fundamental programmes of cellular resilience rather than blocking individual death pathways.

Transcriptomic profiling revealed the molecular basis of this broad cytoprotection. Y4R activation drives coordinated transcriptional reprogramming that enhances β-cell identity programmes, with upstream regulator analysis predicting activation of key transcription factors including NEUROD1, SIX2, and HMGA1. Multiple stress defence mechanisms were implicated: predicted activation of the KEAP1-NFE2L2 antioxidant pathway, enhanced proteostatic capacity through SYVN1 and the ubiquitin-proteasome system, and improved mitochondrial quality control via OMA1. A notable feature of this transcriptional signature was the suppression of global translation pathways, including EIF2 signalling and rRNA processing, concurrent with predicted activation of selective translation regulators LARP1 and EIF4E. This pattern is consistent with a shift from bulk protein synthesis toward selective translation of stress-adaptive transcripts, a mechanism that would reduce ER burden, a critical vulnerability of β-cells, whilst maintaining synthesis of essential survival proteins, though direct translational profiling will be required to confirm this interpretation [31–33]. The coordinate suppression of stress-responsive transcription factors including NR4A1 and MLXIPL is consistent with reduced cellular stress burden in K22-treated islets [34, 35]. Collectively, these findings suggest that Y4R activation promotes a state of enhanced β-cell fitness characterised by reinforced identity, selective protein synthesis, and multi-layered stress defences.

Beyond intrinsic cytoprotection, Y4R activation exerts immunomodulatory effects within the islet microenvironment. The coordinated suppression of chemokines that recruit pathogenic immune cells, alongside preservation of IL-2 and Foxp3 expression, suggests that K22 remodels the local immune landscape toward a less pro-recruitment, more regulation-supportive environment. In contrast, Y4R agonism appears to operate at the site of autoimmune attack, reducing the signals that draw effector cells to islets whilst maintaining pro-regulatory markers locally. Critically, this immunomodulatory effect extends to a fully human system. K22 significantly attenuated IL-2-activated, T cell-enriched human PBMC migration toward cytokine-stressed human islets across multiple timepoints, demonstrating conservation of the anti-recruitment mechanism in human immune-islet interactions. This dual action, enhancing β-cell resilience whilst dampening local immune recruitment, is consistent with the observed delay in diabetes onset in vivo.

The demonstration that Y4R agonism delays autoimmune diabetes onset provides critical validation of the therapeutic potential suggested by our mechanistic studies. The adoptive transfer model employed, using diabetogenic NY8.3 CD8⁺ T cells transferred into Rag1-deficient recipients, represents a stringent test of efficacy against a defined autoreactive T cell population [37, 38]. Disease delay in this setting, where endogenous adaptive immune regulation is absent, indicates that K22 acts through mechanisms other than enhancement of systemic tolerance. The altered CD8⁺ T cell dynamics observed in pancreatic lymph nodes, increased accumulation with reduced naïve phenotype, are consistent with altered trafficking or retention of effector cells [39], aligning with our in vitro findings of reduced chemokine production and impaired T cell migration toward K22-treated islets. Whilst the precise contribution of intrinsic β-cell protection versus local immunomodulation to the observed disease delay cannot be definitively separated, the convergence of both mechanisms at the islet level likely underlies the therapeutic effect. This positions Y4R agonism as a candidate first-in-class approach that addresses both the target of autoimmune attack and the local inflammatory milieu simultaneously.

The dual mechanism of Y4R agonism, combining intrinsic β-cell protection with local immune modulation, suggests several potential clinical applications beyond early-stage T1D. Islet transplantation remains limited by early graft loss due to inflammatory stress and immune-mediated destruction, and the requirement for chronic immunosuppression restricts broader adoption [40]. An agent that enhances graft resilience whilst reducing local immune attack could improve engraftment efficiency and potentially reduce immunosuppressive burden. Similar considerations apply to emerging stem cell-derived β-cell therapies, where protecting transplanted cells from both inflammatory damage and immune rejection remains a critical challenge [41, 42]. Evidence that K22 protects human islets from inflammatory damage and reduces human immune cell recruitment toward human islets provides a strong rationale for testing Y4R agonism in preclinical islet transplantation and stem cell-derived β-cell therapy settings. Pairing local islet protection with systemic immune modulation, such as anti-CD3 therapy, could potentially achieve greater efficacy or durability than either approach alone, whilst allowing dose reduction of systemic immunosuppressants.

Our findings raise broader questions about the physiological significance of endogenous PP/Y4R signalling in islet resilience. β-cells are intrinsically susceptible to oxidative and biosynthetic stress due to their unusually low antioxidant defences and high ER secretory demand, making tonic cytoprotective signalling physiologically plausible. That K22 reduces basal caspase-3/7 activity and suppresses NR4A1 without exogenous stressors suggests Y4R engagement provides constitutive buffering against this vulnerability. The preservation of GSIS and whole-body glucose tolerance in K22-treated mice further supports a stress-conditional rather than nutrient-sensing role for Y4R, consistent with the absence of reported islet phenotypes in Y4R knockout models under homeostatic conditions and synergistic metabolic effects only in Y2Y4 double knockouts. Furthermore, a DPP-4-resistant PP analogue preserves β-cell area and reduces apoptosis in obesity-associated diabetes [21, 43], aligning with our cytoprotective findings across distinct diabetogenic contexts. Whether progressive loss of PP signalling during insulitis contributes to β-cell vulnerability by withdrawing tonic oxidative protection and chemokine restraint simultaneously remains a compelling and testable question.

Several limitations warrant consideration. The transcriptomic profiling was performed under basal culture conditions, and whether the identified resilience programme operates equivalently under active inflammatory challenge remains to be directly tested. The adoptive transfer model employed does not fully capture the complexity of spontaneous autoimmune diabetes, and the dosing regimen was not pharmacokinetically optimised, suggesting that the observed disease delay likely represents a conservative estimate of therapeutic potential. For translation, Y4R is expressed beyond islets, including CNS satiety circuits and the gastrointestinal tract, raising the possibility of off-target extra-pancreatic effects. Tissue distribution, CNS exposure, tolerability, and longer-term safety will therefore be key parameters for clinical development. The human PBMC migration data were obtained using IL-2-activated, non-HLA matched donors, reflecting cytokine-driven rather than antigen-specific T cell recruitment, and whether K22 suppresses antigen-specific human T cell recruitment to islets remains to be determined.

In conclusion, we identify Y4R as a promising therapeutic target for type 1 diabetes that operates through a fundamentally different mechanism to current immunomodulatory approaches. The combination of direct β-cell cytoprotection, local immune modulation, and demonstrated *in vivo* efficacy establishes Y4R agonism as a candidate strategy for disease modification. By enhancing intrinsic β-cell resilience whilst dampening local immune attack, Y4R targeting may offer therapeutic benefit without the risks of systemic immunosuppression, though this will require validation through dedicated pharmacokinetic and long-term safety studies. These findings open new avenues for preserving endogenous β-cell function in T1D, enhancing graft survival in islet transplantation, and protecting stem cell-derived β-cells from inflammatory damage.

## Acknowledgements

Human islets were obtained from the King’s College London Human Islet Research Tissue Bank (KCL HI-RTB; 20/SW/0074). We are deeply grateful to the families of pancreas donors, as well as to the Cell Therapy Unit within the NIHR/Wellcome King’s Clinical Research Facility, which hosts the clinical islet isolation laboratory, for their vital support in providing islets for this study. We would also like to thank Dr. Niwa Ali and Dr. Prudence Lui for providing the Foxp3 transgenic mice.

## Author contributions

G.A.B conceived and designed the study and wrote the manuscript. N.A.H. planned and performed experiments and contributed to manuscript writing. K.W.T. and E.O.O. performed experiments and assisted with manuscript editing. N.H.F.F contributed to imaging of islets. M.Z, Y.L, MH and M.K.M.M. contributed to manuscript editing. R.M synthesized the K22 peptide. J.A.P designed and performed the in vivo experiments and edited the manuscript. A.I analysed the flow cytometry data. S.J.P., D.J.H, A.G.B-S and G.A.B. provided critical review and edited the manuscript.

## Data availability

The RNA-sequencing data that support the findings of this study are available in the Gene Expression Omnibus (GEO) under accession number GSE 331080. All other data supporting the conclusions of this article are available from the corresponding author upon reasonable request.

## Methods

### Experimental models

#### 2.1 Animals

Male CD-1, male/female C57BL6/J and male BALB/c mice were obtained from Charles River Laboratories and maintained under specific pathogen-free conditions at 22 ± 2 °C on a 12-h light/dark cycle with ad libitum access to standard chow and water. Recombinase-activating gene–deficient NOD (Rag1⁻^/^⁻ NOD), NY8.3 T cell receptor–transgenic (TCR-Tg) NOD, and BDC2.5 TCR-Tg NOD mice were originally sourced from The Jackson Laboratory. These strains have been maintained in-house at Cardiff University, where *in vivo* experiments for this study were performed by Dr James Pearson. Splenocytes for CD8⁺ T cell isolation was obtained from four 8-week-old female diabetic NY8.3 NOD mice (a generous gift from Dr James Pearson, Cardiff University). For experiments evaluating Foxp3 expression, Foxp3-GFP-DTR C57BL/6 male mice were used, which were a kind gift from Dr Niwa Ali, King’s College London. All animal experiments were conducted under a UK Home Office project licence and in accordance with the Animals (Scientific Procedures) Act 1986.

#### 2.2 Human pancreatic islets

Human pancreatic islets were obtained from non-diabetic, heart-beating, brain-dead organ donors and isolated by cold collagenase digestion as previously described (21, 61), under ethical approval from the King’s College London Human Islet Research Tissue Bank (KCL HI-RTB; 20/SW/0074). Islets from three donors were used: donor 1, a 41-year-old female (BMI 24), donor 2, a 36-year-old male (BMI 23) and donor 3, a 27-year-old male with a BMI of 24.7.

#### 2.3 Mouse islet isolation and culture

Pancreatic islets were isolated from male CD-1 or male/female C57BL6/J mice (8–12 weeks old) by collagenase digestion of the exocrine pancreas followed by density and size-based separation of intact islets. After washing to remove residual exocrine tissue and enzyme, islets were hand-picked under a stereomicroscope to ensure high purity. Purified mouse islets were then recovered overnight at 37 °C in a humidified incubator (95% air/5% CO₂) in RPMI-1640 medium containing 10% (v/v) foetal bovine serum, 100 U/ml penicillin and 100 µg/ml streptomycin before experimental use.

### Key reagents and interventions

#### 2.4 K22 synthesis and formulation

The Y4R agonist K22 and its fluorescent analogue K22-TAMRA have been synthesized as described previously [1]. K22, previously named compound 4b or [K22(PEG22)] hPP2-36 [1] consists of hPP bearing a site-specific introduction of Lys at position 22 modified selectively with a ca 22 kDa polyethylene glycol (PEG) moiety by a stable linker, which confers enhanced plasma stability and prolonged *in vivo* activity while preserving high selectivity for NPY4R. The fluorescent tracer K22–TAMRA (TAMRA-4b) corresponds to the same PEGylated hPP scaffold bearing an additional 5-carboxytetramethylrhodamine (TAMRA) fluorophore at the N-terminus for imaging applications. Purity of the peptides has been confirmed by HPLC and mass spectrometry.

For all *in vitro* experiments, lyophilised K22 and K22–TAMRA were reconstituted in sterile water to 12 µM stock solutions, aliquoted, and stored at –80 °C to avoid repeated freeze–thaw cycles. Working concentrations (6–500 nM) were prepared immediately before use by dilution into cell culture medium containing 10% FBS and 1% Penicillin and streptomycin. For *in vivo* studies, K22 was diluted in sterile 0.9% saline and administered by subcutaneous injection at the indicated doses and schedules.

#### 2.5 Antibodies, fluorescent probes and other reagents

Primary and secondary antibodies for immunofluorescence and flow cytometry were selected to identify endocrine cell subsets, proliferating and apoptotic β cells, and leukocyte populations, and were used at the same concentrations and under similar conditions as in reformed islet preparations. Primary antibodies included reagents directed against insulin, glucagon, somatostatin, pancreatic polypeptide, Ki67, cleaved caspase-3, CD45, CD8, CD4 and F4/80, with species-specific secondary antibodies conjugated to Alexa Fluor dyes for multiplex imaging; full antibody details are provided in Supplementary Tables 1 and 2. Nuclear staining used DAPI, while live/dead discrimination employed amine-reactive viability dyes, and caspase-3/7 activity was quantified with the Caspase-Glo 3/7 Assay Kit (Promega). Pro-inflammatory cytokines for diabetogenic stress (recombinant murine TNF-α, IFN-γ and IL-1β) were obtained from PeproTech, and additional stressors, including streptozotocin, palmitate and thapsigargin, as well as collagenase type XI, Histopaque-1077, culture media and supplements, were purchased from Sigma-Aldrich. CD8⁺ T cells were isolated from spleens of four 8-week-old diabetic female NY8.3 NOD mice using a MojoSort Mouse CD8 T-cell isolation kit and MojoSort buffer (BioLegend), and T-cell migration and chemotaxis were assessed using CytoSelect and EZCell cell migration/chemotaxis assay kits (Cell Biolabs and BioVision, respectively).

**Table 1.** Primary Antibodies.

| Target | Company | Product # | Host species | Concentration |
| --- | --- | --- | --- | --- |
| Insulin | Agilent | IR00261-2 | Guinea pig | 1:10 |
| Glucagon | Merck Life Science UK Limited | G2654 | Rabbit | 1:1000 |
| Somatostatin | Abcam | ab30788-125ul | Rabbit | 1:50 |
| Ki67 | Abcam | ab15580 | Rabbit | 1:200 |
| CD80 | Bio-Techne | AF740-SP | Goat | 1:100 |
| IBA1 | Alpha Laboratories Limited | 019-19741 | Rabbit | 1:200 |
| CD45 | Abcam | ab40763 | Rabbit | 1:100 |
| CD8α | Cell Signaling | 98941S | Rat | 1:100 |
| CD4 | BD Pharmingen™ | 557667 | Rat | 1:50 |
| Cleaved caspase-3 | Cell Signaling | 9661S | Rabbit | 1:100 |

**Table 2.** Secondary Antibodies.

| Target | Company | Product # | Concentration |
| --- | --- | --- | --- |
| Alexa Fluor® 488-AffiniPure Donkey Anti-Guinea Pig IgG | Strattech | 706-545-148-JIR | 1:200 |
| Donkey anti-Rabbit IgG (H+L) Highly Cross-Adsorbed Secondary Antibody, Alexa Fluor™ 568 | Invitrogen | A10042 | 1:1000 |
| Donkey anti-Mouse IgG (H+L) Highly Cross-Adsorbed Secondary Antibody, Alexa Fluor™ Plus 647 | Invitrogen | A32787 | 1:1000 |
| Donkey anti-Rat IgG (H+L) Highly Cross-Adsorbed Secondary Antibody, Alexa Fluor™ Plus 647 | Invitrogen | A48272 | 1:1000 |
| Donkey anti-Goat IgG (H+L) Cross-Adsorbed Secondary Antibody, Alexa Fluor™ 594 | Invitrogen | A-11058 | 1:1000 |

### In vitro functional assays

#### 2.6 Insulin secretion from mouse and human reformed islets

Insulin secretion from native mouse and human islets was assessed using static incubation assays. Islets were first pre-incubated for 1 h in physiological salt solution (Gey & Gey, 1936) containing 2 mM glucose. Subsequently, groups of five islets per replicate (n = 6–10 for each condition) were incubated in appropriate volumes (100 µl buffer per islet) of Gey & Gey solution supplemented with either 2 mM or 20 mM glucose. Following incubation, supernatants were collected and stored at −20 °C until analysis of insulin concentration by radioimmunoassay.

#### 2.7 Dynamic perifusion insulin secretion and calcium imaging

Dynamic glucose-stimulated insulin secretion was assessed using a perifusion system. Batches of size-matched mouse islets were loaded into perifusion chambers and equilibrated in Krebs–Ringer bicarbonate buffer containing 2 mM glucose, then sequentially exposed to low and high glucose solutions (e.g. 2 mM then 20 mM glucose) at a constant flow rate; effluent fractions were collected at 2 mins intervals and insulin concentrations were measured by radioimmunoassay. In parallel, single-islet calcium imaging was performed by loading islets with a fluorescent Ca²⁺ indicator, Fura-2, followed by time-lapse imaging during sequential low and high glucose and ATP stimulation; traces were analysed to quantify glucose-evoked Ca²⁺ influx and network connectivity/hub-cell organisation.

#### 2.8 Islet connectivity imaging and analysis

Islets were loaded with Fluo-8 and imaged using a Crest X-Light spinning-disk confocal system, Nikon Ti-E microscope and 10×/0.4 NA air objective (MetaMorph v7.10.3; Molecular Devices). Fluo-8 was excited at 458–482 nm using a Lumencor Spectra X light engine, and fluorescence emission was collected at 500–550 nm using a Photometrics Delta Evolve EM-CCD camera. Imaging was performed in HEPES-bicarbonate buffer containing (mmol/L): 120 NaCl, 4.8 KCl, 24 NaHCO₃, 0.5 Na₂HPO₄, 5 HEPES, 2.5 CaCl₂, 1.2 MgCl₂, and 3–17 D-glucose.

Functional connectivity analysis and hub cell identification were performed using an in-house matrix binarization pipeline as previously described [10.1038/s41467-020-20632-z and 10.1016/j.cmet.2016.06.020]. Individual cell Ca²⁺ traces were extracted from manually defined ROIs, processed, binarized, and subjected to permutation-based connectivity analysis. Hub cells were defined as cells exhibiting 60–100% of the significant functional connections.

#### 2.9 Apoptosis: Caspase-Glo 3/7 assay

Apoptosis was quantified using the Caspase-Glo 3/7 assay (Promega) to measure caspase-3/7 activity under four experimental conditions: (1) untreated control, (2) K22-treated, (3) cytokine-treated during the final 24 h, and (4) K22-pretreated followed by combined K22 plus cytokine treatment during the final 24 h (K22 + CK). Primary islets were cultured in RPMI medium supplemented with 2% FBS and 1% penicillin/streptomycin and, where indicated, pretreated with K22 at 12nM for 24 h before cytokine exposure. Cytokine conditions consisted of a pro-inflammatory cocktail (CK) containing IL-1β (0.05 U/µl), TNF-α (1 U/µl), and IFN-γ (1 U/µl). Caspase-3/7 activity was then determined by bioluminescence using the Caspase-Glo 3/7 assay in accordance with the manufacturer’s instructions.

#### 2.10 RNA Extraction and Real time qPCR

Total RNA was extracted from isolated mouse and human islets using TRIzol reagent according to the manufacturer’s instructions. Briefly, islets were lysed in TRIzol, phase-separated with chloroform, and the aqueous phase was collected for RNA precipitation with isopropanol, followed by ethanol washing and resuspension of the RNA pellet in nuclease-free water. RNA concentration and purity were assessed spectrophotometrically, and samples with acceptable A260/280 ratios were used for downstream applications.

For cDNA synthesis, 500–1,000 ng of total RNA per sample was reverse transcribed using a high-capacity cDNA reverse transcription kit with random primers, following the manufacturer’s protocol. Quantitative PCR (qPCR) was performed using TaqMan Gene Expression Assays on a real-time PCR system in 96-well plate format, with each reaction containing TaqMan Universal Master Mix, gene-specific TaqMan probe/primer sets, and diluted cDNA in a total volume of 10–20 µl. Reactions were run in technical triplicates, and relative gene expression was calculated using the ΔΔCt method with a housekeeping gene (Mm03928990_g1) as internal control; data are presented as delta CT or fold-change relative to untreated controls.

##### 2.10.1 Foxp3 expression in Foxp3–GFP–DTR islets treated with K22, cytokines and diphtheria toxin

Pancreatic islets were isolated from Foxp3–GFP–DTR C57BL/6 male mice (a kind gift from Dr Niwa Ali) as described above and recovered overnight at 37 °C in RPMI-1640 supplemented with 10% (v/v) FBS, 1% penicillin/streptomycin, and 2 mM L-glutamine. The following day, islets were allocated to the indicated treatment conditions: RPMI control, cytokine cocktail (CK), K22 plus cytokines (K22+CK), and, where specified, diphtheria toxin (DT) to deplete Foxp3–GFP–DTR⁺ cells. K22 was used at 12 nM, and the cytokine cocktail contained IL-1β, TNF-α and IFN-γ at the concentrations described above. DT was added at 50 ng/ml from a 1,000 ng/µl stock and cultures were maintained for 24 h at 37 °C in a humidified 5% CO₂ incubator.

At the end of the culture period, islets were washed twice in ice-cold PBS and lysed directly in TRIzol reagent. Total RNA was extracted according as described above. For cDNA synthesis, 500–1,000 ng of total RNA per sample was reverse-transcribed using a high-capacity cDNA reverse transcription kit with random primers, following the manufacturer’s protocol. Quantitative PCR (qPCR) was performed using TaqMan Gene Expression Assays and TaqMan probe for Foxp3. Reactions were run in technical triplicates. Relative Foxp3 expression was calculated using the ΔΔCt method with the housekeeping gene as internal control, and data were expressed as -ΔCt, with DT-treated samples serving as a functional control for Foxp3 ablation, as shown in Figure 5M.

#### 2.11 Multiplex cytokine/chemokine assays and CXCL10 ELISA

To characterise the islet inflammatory milieu, cytokine and chemokine levels were measured in supernatants or lysates from mouse islets cultured under control, K22, cytokine, and K22 + cytokine conditions. CXCL10 concentrations were quantified using a specific ELISA according to the manufacturer’s protocol, while broader panels of analytes (including CCL2, CCL3, CCL4, CCL7, CXCL2, IL-6, IL-2 and others) were measured using multiplex bead-based immunoassays. Data were log-transformed where appropriate, non-detects handled as per kit guidance, and values normalised to protein content before statistical analysis.

#### 2.12 NY8.3 CD8⁺ T-cell activation and proliferation assays

NY8.3 CD8⁺ T cells were isolated from spleens of NY8.3 TCR-Tg NOD mice and cultured with irradiated splenic antigen-presenting cells pulsed with the cognate IGRP₂₀₆–₂₁₄ peptide. Cells were stimulated across a range of peptide concentrations in the presence or absence of K22, and proliferation was assessed after culture (e.g. by CFSE dilution or thymidine/EdU incorporation), while supernatants were collected for measurement of effector cytokines (IFN-γ, TNF-α, MIP-1β) by ELISA. Npy4r transcript levels in resting and activated NY8.3 cells were quantified by qPCR to assess induction of this pathway upon antigen stimulation.

### Immune–islet interaction assays and imaging

#### 2.13 Immunofluorescence and confocal microscopy

Immunofluorescence staining was performed to localize Y4R receptor expression for RNAscope and K22-TAMRA studies, as well as to assess cell proliferation, K22-mediated protection of cytokine-damaged islets, and in various invasion assays. Whole native islets were fixed in 4% paraformaldehyde (PFA) and subsequently incubated in blocking buffer (1% BSA and 10% donkey serum in 0.1% PBST) for 2 h at room temperature to prevent non-specific antibody binding. The islets were then incubated overnight at 4 °C with primary antibodies against insulin, glucagon, somatostatin, Ki67, CD80, IBA1, CD8, CD3e, and CD4 (see Supplementary Table 1), followed by incubation with appropriate fluorophore-conjugated secondary antibodies for 1 h at room temperature (see Supplementary Table 2). Nuclei were counterstained and mounted using DAPI Fluoromount-G. To confirm antibody specificity, primary and secondary antibody control experiments were carried out to exclude any endogenous background staining for insulin and CD8. Fluorescence images of islets were acquired using an Eclipse Ti-E Inverted A1 or a Zeiss LSM700 confocal microscope and analysed with CellProfiler and the Cell Counter plugin in ImageJ software.

#### 2.14 Islet destruction: Migration assay

The Transwell migration assay was used to evaluate the chemotactic response of immune cells toward islet-derived signals and to examine the impact of K22 under inflammatory conditions. Islets were isolated from CD1 mice, BALB/c mice, and human cadaveric donors, with BALB/c mice selected because they express the H2Kᵈ MHC class I haplotype, providing compatibility with the antigen-specific T cells used in these experiments. The islets were cultured in RPMI medium supplemented with 2% FBS and 1% penicillin/streptomycin for 24 h, after which K22 was employed in selected migration and invasion assays.

In parallel, RAW 264.7 macrophage-like cells were activated with LPS (200 ng/ml) and IFN-γ (2.5 ng/ml) for 24 h. After activation, cells were washed with PBS to remove residual LPS and IFN-γ, detached by trypsinisation, pelleted, and 1 × 10⁶ cells were seeded into the inserts (upper chamber) placed above coverslips containing 3–4 reformed islets for a further 24 h. CD8⁺ T cells were isolated from the spleens of four 8-week-old female NY8.3 NOD mice; a single-cell suspension was prepared in MojoSort buffer at 1 × 10⁸ cells/ml, incubated with a biotinylated antibody cocktail (anti-CD4, CD11b, CD11c, CD19, CD24, CD45R/B220, CD49b, CD105, I-A/I-E, TER-119/erythroid, and TCRγδ) followed by streptavidin nanobeads, and magnetically separated, with the unbound fraction collected as enriched CD8⁺ T cells.

Similarly, CD4^+^ T cells were isolated from the spleens of four 8-week-old female BDC NOD mice using MojoSort™ Mouse CD4 Naïve T Cell Isolation Kit (cat. No.480040) and following manufacturer’s instruction.

Peritoneal macrophages were also obtained as an additional primary macrophage source. Briefly, 10 ml sterile PBS was injected into the peritoneal cavity while gently lifting the abdominal wall to form a tent, directing the fluid into the peritoneal fat pad. The abdomen was gently massaged for 10–25 s, and the peritoneal lavage fluid was slowly aspirated with a fresh 10 ml syringe fitted with a 25-gauge needle, taking care to avoid clogging by fat or organs. The lavage was collected into a 50 ml conical tube kept on ice and centrifuged at 300–500 × g for 10 min at 4 °C to pellet the cells; the pellet was resuspended in cold RPMI and cultured at 37 °C in a 5% CO₂ incubator to allow macrophage adherence, after which non-adherent cells were removed by gentle PBS washes, yielding a highly enriched peritoneal macrophage population (typically >90%).

For CD8⁺ T-cell migration assays, an 8 µm-pore Transwell plate (BioVision, Inc., USA) was used according to the manufacturer’s instructions. BALB/c reformed islets were cultured for 48 h in complete RPMI containing 2% FBS in the lower chamber, and 2 × 10⁵ CD8⁺ T cells were seeded into the upper chamber positioned over coverslips bearing reformed islets that were either untreated or exposed to a pro-inflammatory cytokine cocktail. After 18 h, cells that had migrated to the lower chamber were quantified by measuring optical density at Ex/Em = 540/590 nm using a PHERAstar microplate reader.

To examine the effect of K22 on macrophage migration and islet destruction in the presence of pro-inflammatory cytokines, similar Transwell-based assays were performed using either RAW 264.7 cells or primary peritoneal macrophages. Mouse or human islets were incubated for 48 h in complete RPMI with 2% FBS, with or without K22 (12 nM) during the first 24 h. Following this pretreatment, activated RAW 264.7 cells or peritoneal macrophages (2 × 10⁵ cells) were seeded into the upper chamber and placed over coverslips with reformed islets; pro-inflammatory cytokines were added for the final 24 h. After incubation, cells that had migrated into the lower chamber were quantified by measuring O.D. values at Ex/Em = 540/590 nm using a PHERAstar microplate reader.

#### 2.15 Islet destruction: Invasion assay

A Transwell-based invasion assay was established to evaluate both chemotactic responses and the ability of immune cells to infiltrate islets through an extracellular matrix, a strategy commonly applied in studies of cancer metastasis and embryonic development. Using this approach, the impact of cytokine-induced islet damage in the presence or absence of K22 was examined in mouse and human islets co-cultured with CD8⁺ T cells, activated RAW 264.7 monocyte/macrophage-like cells, and peritoneal macrophages, with outcomes assessed by confocal imaging.

For these experiments, coverslips with adhered native islets recovered from the migration assays described above were used. The coverslips were fixed and immunostained with antibodies against insulin, CD8, or CD80, followed by DAPI nuclear counterstaining, and reformed islet morphology and immune cell infiltration were qualitatively evaluated by light and confocal microscopy.

#### 2.16 Migration and invasion assays using human PBMCs and human pancreatic islets

Human pancreatic islets were obtained and recovered as described above. Following isolation, islets were briefly identified by dithizone staining using a minimalist quick-stain protocol, hand-picked into dithizone-free medium, and allowed to recover for 18–24 h in supplemented CMRL-1066 containing FBS (10% v/v), B-27 supplement (1×), HEPES (10 mM), GlutaMAX (2 mM), penicillin/streptomycin (1%), nicotinamide (10 mM), and glucose (11 mM) before use in migration and invasion experiments. Islets were then transferred onto Matrigel-coated coverslips and cultured at 37 °C in a humidified 5% CO₂ incubator.

Cryopreserved human peripheral blood mononuclear cells (PBMCs) were rapidly thawed at 37 °C, diluted dropwise into warm complete RPMI-1640 supplemented with FBS (10% v/v) and penicillin/streptomycin (1%), washed twice at 300 × g for 10 min to remove residual DMSO, and counted by trypan blue exclusion. PBMCs were resuspended at 1 × 10⁶ cells/ml in complete medium and expanded for 2 days with CD3/CD28 beads at a 1:1 bead-to-cell ratio prior to assay. PBMCs were further activated with IL-2 (50 U/L). For imaging experiments, cells were labelled with Qtracker 525 according to the manufacturer’s instructions by incubating cells with pre-complexed labelling reagent for 45–60 min at 37 °C protected from light, followed by two washes in complete RPMI.

For treatment conditions, human islets were allocated to four groups: vehicle (RPMI containing 2% FBS and 1% penicillin/streptomycin), K22, cytokine cocktail (CK), or K22 plus cytokine cocktail (K22+CK). Islets were pretreated for 24 h with K22 (12 nM) or vehicle, after which an inflammatory challenge was applied for a further 24 h where indicated using a cytokine cocktail containing IL-1β (0.05 U/µl), TNF-α (1 U/µl), and IFN-γ (1 U/µl).

Transwell migration assays were performed using inserts placed above wells containing human islets in the lower chamber. For each condition, 10–15 human islets were added to 600 µl medium in the lower chamber, with or without K22 and/or cytokines as described above. PBMCs were resuspended in serum-free medium and seeded into the upper chamber at approximately 3 × 10⁵ cells in 300 µl per insert. Plates were incubated at 37 °C and 5% CO₂ for 2–20 h, after which inserts were removed and the medium from the lower chamber was collected to quantify migrated cells.

Migrated PBMCs were pelleted by centrifugation at 1,000 × g for 5 min, washed in assay wash buffer, and incubated with dissociation solution/cell dye mixture according to the migration assay kit instructions. Fluorescence was measured on a plate reader at Ex/Em 530/590 nm, and background-subtracted fluorescence values were converted to cell numbers using a standard curve generated in parallel from serial dilutions of PBMCs processed under identical dye-loading conditions. Migration data were expressed as the total number of migrated cells per well.

For invasion assays, coverslips containing human islets recovered from Transwell co-cultures were fixed in 4% paraformaldehyde and processed for immunofluorescence as described above. Islets were stained with antibodies against CD45 and insulin and counterstained with DAPI, then imaged by confocal microscopy to assess PBMC infiltration into islet structures and associated disruption of islet architecture. Invasion of PBMCs was additionally visualised by direct detection of Qtracker-labelled immune cells within or around the islet mantle. Images were analysed qualitatively to compare the extent of immune cell invasion across treatment groups.

#### 2.17 DA-ZP1-based sorting of β-cell–enriched islet fractions

Mouse pancreatic islets were isolated from 10-week-old male CD1 mice and recovered overnight in RPMI 1640 supplemented with 10% (v/v) FBS and 1% (v/v) penicillin–streptomycin at 37 °C in a humidified 5% CO₂ incubator. The following day, intact islets were stained with the zinc-reactive probe DA-ZP1 by adding 5 µl of a 20 mM DA-ZP1 stock solution to 5 ml of culture medium and incubating for 30 min at 37 °C on an orbital rocker. Islets were then washed twice in FACS buffer (PBS containing 2% FBS and 2 mM EDTA). Approximately 500 islets were transferred into each of three microcentrifuge tubes and dissociated with TrypLE Express at 37 °C for 15 min in a dry-block incubator, followed by gentle trituration to obtain single-cell suspensions. Cells were filtered through a 40 µm strainer and subjected to flow-cytometric sorting based on DA-ZP1 fluorescence and 7-AAD viability staining; DA-ZP1⁺ (β-cell–enriched) and DA-ZP1⁻ fractions of live single cells were collected separately for downstream analyses.

#### 2.18 RNAscope in situ hybridisation for Npy4r

RNAscope in situ hybridisation was performed on mouse pancreatic sections to localise Npy4r transcripts within islets. Formalin-fixed paraffin-embedded sections were processed using the RNAscope Multiplex Fluorescent assay (ACD Bio) according to the manufacturer’s instructions, with probes specific for Npy4r alongside positive (Ubc) and negative (DapB) controls, followed by immunostaining for insulin and nuclear counterstaining with DAPI. Fluorescent puncta corresponding to Npy4r mRNA were imaged by confocal microscopy, and signal density within insulin-positive regions was quantified using ImageJ / CellProfiler to assess beta-cell–restricted expression.

#### 2.19 Fluorescent K22–TAMRA binding and competition

To assess Y4R accessibility at the β-cell surface, fluorescent K22–TAMRA competition binding assays were performed on mouse and human pancreatic sections. Islets were incubated with 12 nM K22–TAMRA in islet culture medium for a 2h in presence and absence of 100-fold concentration of pancreatic polypeptide (PP) at 37 °C. The islets were then washed to remove unbound ligand, and co-immunostained for insulin and/or Nkx6.1 before mounting with DAPI-containing medium. TAMRA signal within insulin-positive areas was quantified by confocal microscopy to confirm specific, PP-blockable binding.

### In vivo studies and downstream analyses

#### 2.20 NY8.3→Rag1⁻/⁻ adoptive transfer model and IPGTT

For *in vivo* efficacy studies, Rag1-deficient NOD mice received *in vitro*–activated NY8.3 CD8⁺ T cells by intravenous adoptive transfer to induce autoimmune diabetes. Briefly, 5×10^6^ splenocytes from NY8.3 TCR-Tg NOD mice were activated with 20ng/ml IGRP₂₀₆–₂₁₄ *in vitro* for 48 hours, CD8^+^ T cells were then isolated by magnetic isolation, and 3×10^6^ were i.v. injected into Rag1⁻/⁻ recipients. Mice were subsequently treated with K22 or vehicle every other day for 13 days. K22 was administered at a dose of 20 mg/kg (20 µg/g, subcutaneously) on each treatment day. Mice were monitored daily for glycosuria from 5 days post-transfer, with blood glucose confirmation using tail vein sampling. Diabetes was defined by sustained hyperglycaemia above 13.9mmol/L (250mg/dl).

#### 2.21 Flow cytometry of islets and lymphoid organs

Single-cell suspensions from mouse islets, spleen, pancreatic draining lymph nodes (PLN), and mesenteric lymph nodes (MLN) were prepared by enzymatic digestion followed by gentle trituration and filtration through 70 µm strainers. Islet preparations were stained with viability dye and fluorochrome-conjugated antibodies against CD45, CD4, CD8, CD11b, CD11c, CD19 and other lineage markers as indicated, to distinguish CD45⁺ immune cells from CD45⁻ endocrine cells and to define lymphoid and myeloid subsets in secondary lymphoid tissues. After staining, cells were washed in FACS buffer (PBS, 2% FBS, 2 mM EDTA) and acquired on a multicolour flow cytometer; data were analysed using standard gating strategies in FlowJo, with doublets excluded by FSC/SSC parameters and gates set using fluorescence-minus-one and isotype controls.

#### 2.22 Bulk RNA-sequencing

Total RNA was isolated from two experimental conditions: untreated islets, and K22-treated islets, with six replicates per condition (200 mouse islets per replicate) using the TRIzol protocol. RNA samples were uniquely barcoded, pooled, and later deconvoluted after sequencing based on their barcode identity.

In vitro transcription was then performed as a linear amplification step to generate amplified RNA (αRNA), which was subsequently fragmented and assessed for integrity using an Agilent TapeStation. The resulting αRNA was reverse transcribed, followed by PCR amplification during which appropriate sequencing adapters were added, yielding a cDNA library that was again quality-checked on an Agilent TapeStation.

Library preparation and bulk RNA-seq were carried out by Single-Cell Discoveries using the QuantSeq 3′ kit (Lexogen), and libraries were sequenced on an Illumina NextSeq 500 to a depth of approximately 10 million reads per sample. Raw reads underwent quality control with FastQC, adapter and poly(A) trimming with BBDuk (BBMap v38.87), and alignment to the mm10 mouse reference genome using STAR/STARsolo (v2.7.10a).

#### 2.23 Data processing

##### 2.23.1 Bulk RNA-Seq analysis

Downstream bioinformatic analysis of the bulk RNA-seq data was performed using the Galaxy platform (usegalaxy.eu). Quality of the raw sequencing reads was first assessed using FastQC, followed by adapter and low-quality sequence trimming with Trimmomatic. Processed reads were then aligned to the appropriate reference genome (mm10 for mouse) using HISAT2, producing BAM files for each sample.

Gene-level read counts were generated using featureCounts (HTSeq-count), quantifying the number of reads mapped to annotated exons based on the corresponding GTF annotation file. The resulting count matrix was imported into DESeq2 within the Galaxy environment for differential gene expression analysis. Samples were assigned to one of two factor levels, untreated and K22-treated. Pairwise comparisons were performed to identify differentially expressed genes (DEGs) with a nominal p-value < 0.05 and a log₂ fold-change threshold as specified. Principal component analysis (PCA) plots and sample-to-sample distance heatmaps were generated from the DESeq2 output to assess sample clustering and replicate reproducibility.

##### 2.23.2 Ingenuity Pathway Analysis

Bulk RNA-seq differential expression results (gene identifiers, log₂ fold changes, and nominal p-values) for the K22-treated versus vehicle-treated comparison were imported into Ingenuity Pathway Analysis (IPA, QIAGEN Inc., Redwood City, CA) for downstream functional interpretation. Genes were filtered prior to upload to retain those with nominal p-value < 0.05 and log₂ fold-change ≥0.5, and the mouse genome was selected as the reference set within IPA.

Core analysis was performed using IPA’s curated knowledge base with default settings, including both direct and indirect relationships, to identify enriched canonical pathways, upstream regulators, and predicted downstream biological effects under basal culture conditions. Canonical pathway analysis identified significantly enriched biological pathways, with activation z-scores used to infer directional activation or inhibition states; pathways with |z-score| ≥ 1.5 and −log(p-value) ≥ 0.5 were considered for interpretation. Upstream regulator analysis predicted the activation states of transcriptional regulators based on the expression patterns of their known downstream target genes, with predictions considered significant at nominal p < 0.05 and |z-score| ≥ 2.0. Causal network and master regulator analyses were performed to identify higher-order regulatory nodes coordinating the observed transcriptional changes, applying the same z-score and p-value thresholds. Results were visualised using IPA’s built-in network and pathway tools. Sequence data are available from the GEO database under accession number GSE331080.

##### 2.23.3 Marker-based cell-type enrichment analysis

To assess the cell-type specificity of K22-induced transcriptional changes, marker-based enrichment analysis was performed using curated gene sets representing the major pancreatic islet endocrine cell types: β-cells (*Ins1*, *Ins2*, *Chga*, *Iapp*, *Nkx6-1*, *Pdx1*, *Ucn3*), α-cells (*Gcg*, *Arx*), δ-cells (*Sst*, *Hhex*), PP cells (*Ppy*), and ε-cells (*Ghrl*). For each gene set, the mean log₂ fold change of constituent genes was calculated from the DESeq2 output and compared against the distribution of log₂ fold changes for all remaining expressed genes using a Wilcoxon rank-sum test. P-values were corrected for multiple testing using the Benjamini–Hochberg method (FDR < 0.05). Enrichment scores are presented as mean log₂ fold change per gene set with associated FDR-corrected p-values.

#### 2.24 Statistical analysis

For murine experiments, approximately 1500-1800 islets were routinely isolated per pancreas, and statistical power calculations were performed for each in vitro assay using data from comparable prior experiments to determine appropriate sample sizes. For 90% power at a 5% significance level to detect differences between treatment groups, static incubation experiments using isolated islets required 4–6 replicates per condition, and islet apoptosis assays required 6 replicates per condition.

Statistical analysis and graphical visualisation were conducted using GraphPad Prism (version 10.5.1; GraphPad Software, La Jolla, CA, USA) and R (v4.5.3). Group comparisons were performed using one- or two-way ANOVA, followed by Šídák’s or Dunnett’s post hoc multiple comparison tests as appropriate. For nested datasets, linear mixed-effects models were used with mouse specified as a random effect. Kaplan–Meier survival curves were analysed using the log-rank (Mantel–Cox) test. All data are presented as mean ± SEM, with statistical significance defined as p < 0.05. Levels of significance are indicated in the figures by asterisks (* p < 0.05; **p < 0.01; *** p < 0.001; **** p < 0.0001), unless otherwise specified.

---

**Figure S1.**
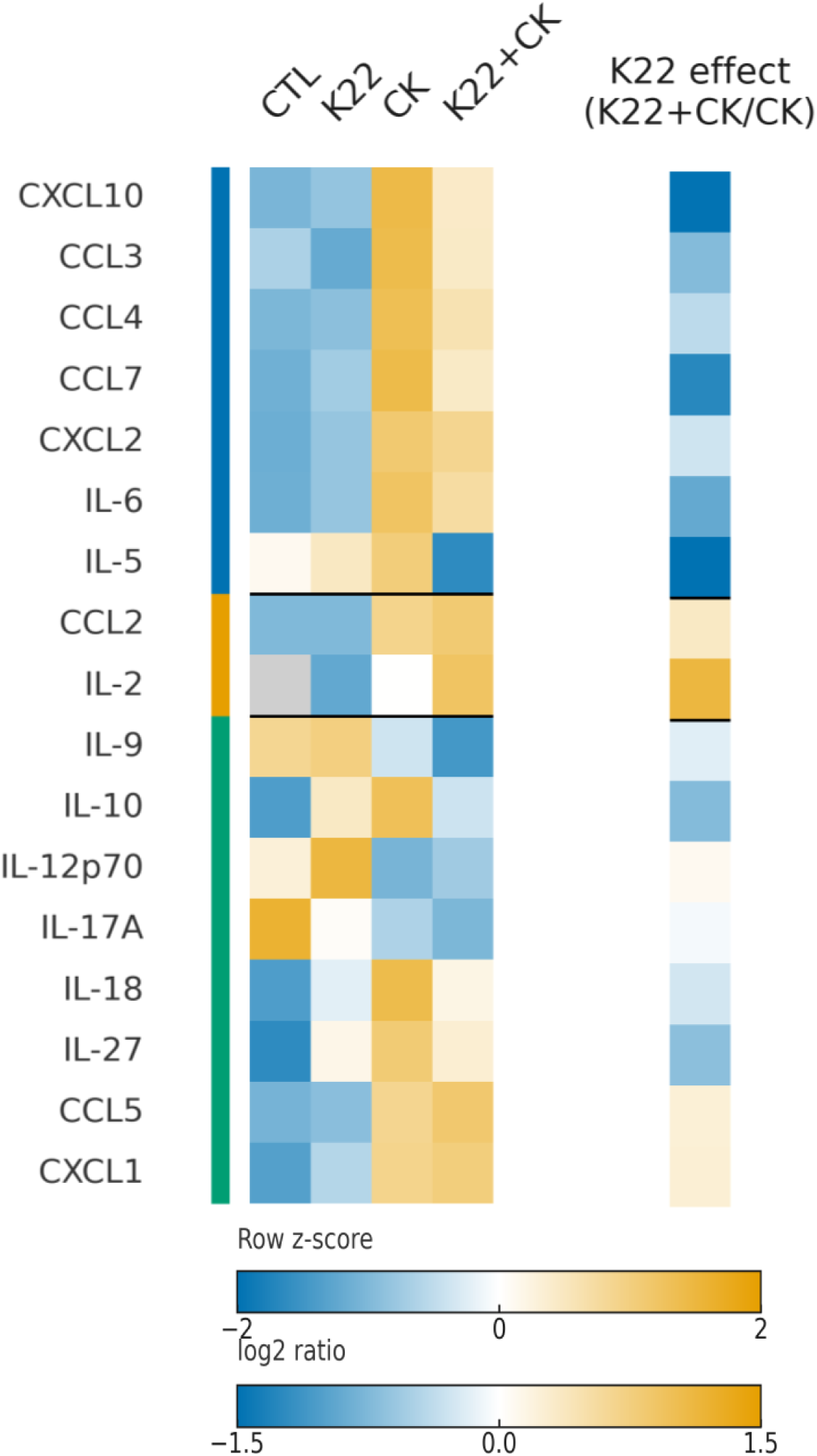
| K22 remodels the cytokine-induced islet inflammatory milieu. Multiplex immunoassay profiling of islet lysates across Control, K22, cytokines (CK), and CK+K22 conditions. The left heatmap shows row z-scores of log10-transformed mean concentrations (pg/mL) for each analyte (non-detects shown in grey), grouped by mediators suppressed by K22, increased by K22, or unaffected. The right heatmap shows the K22 effect in the inflammatory context as log2(K22+CK / CK), where negative values indicate suppression by K22 and positive values indicate enhancement.

**Figure S2.**
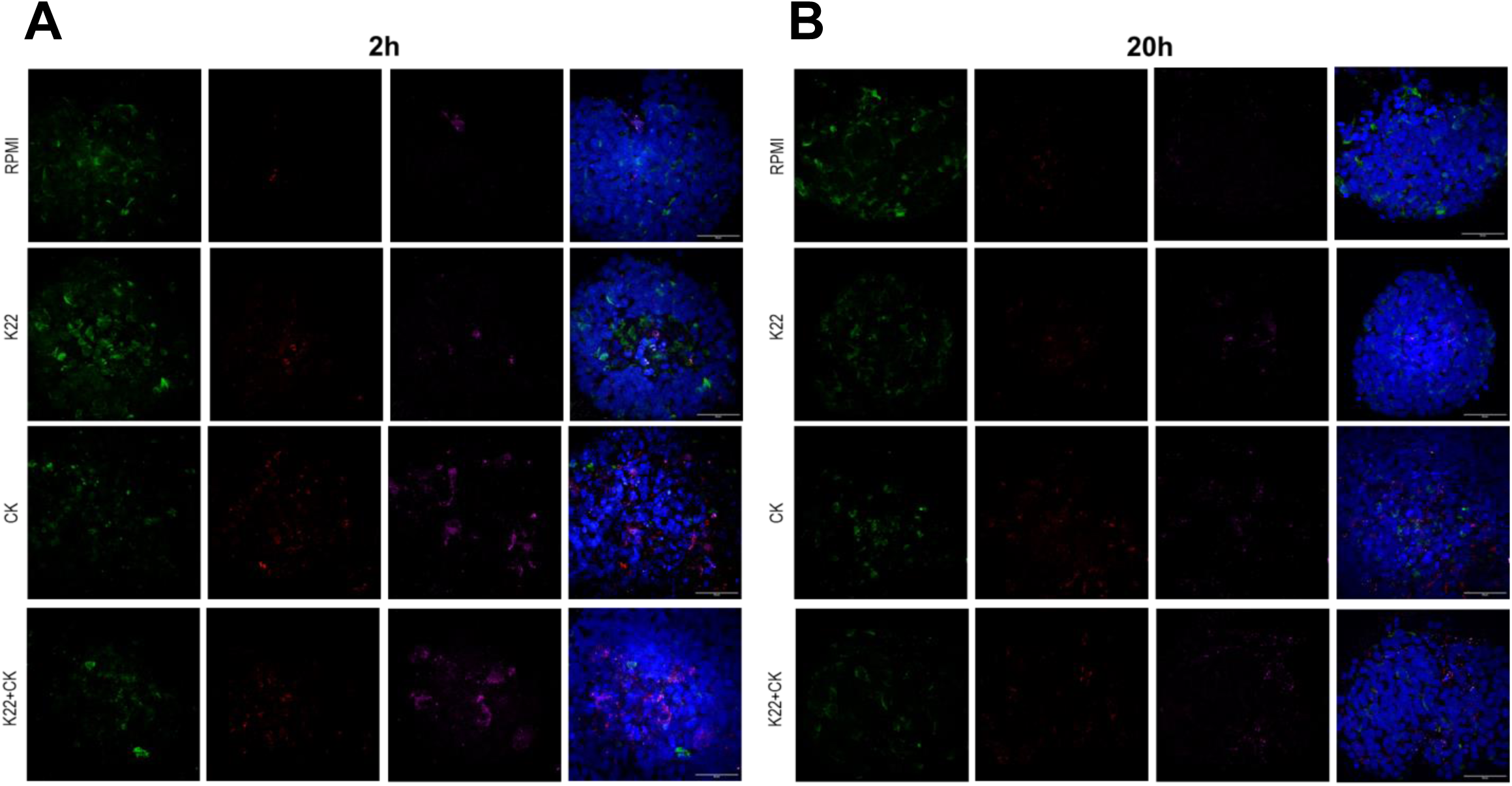
| Time-dependent invasion of activated human PBMCs into human islets. (A) Representative confocal images from a 2 h invasion assay in which activated, Qtracker-labelled human PBMCs were co-cultured with human islets under the indicated conditions (RPMI, K22, cytokines alone (CK), or K22 plus cytokines (K22+CK)). Each row shows Ins (green), Qtracker-labelled PBMCs (red), CD45 (magenta), and the composite image including DAPI (blue), illustrating early PBMC association with the islet surface and increased peri-islet accumulation in the presence of cytokines. (B) Corresponding confocal images acquired after 20 h co-culture for the same conditions as in A, demonstrating progressive PBMC penetration into the islet mantle under cytokine stimulation and reduced invasion when islets are treated with K22 during cytokine exposure.

**Figure S3.**
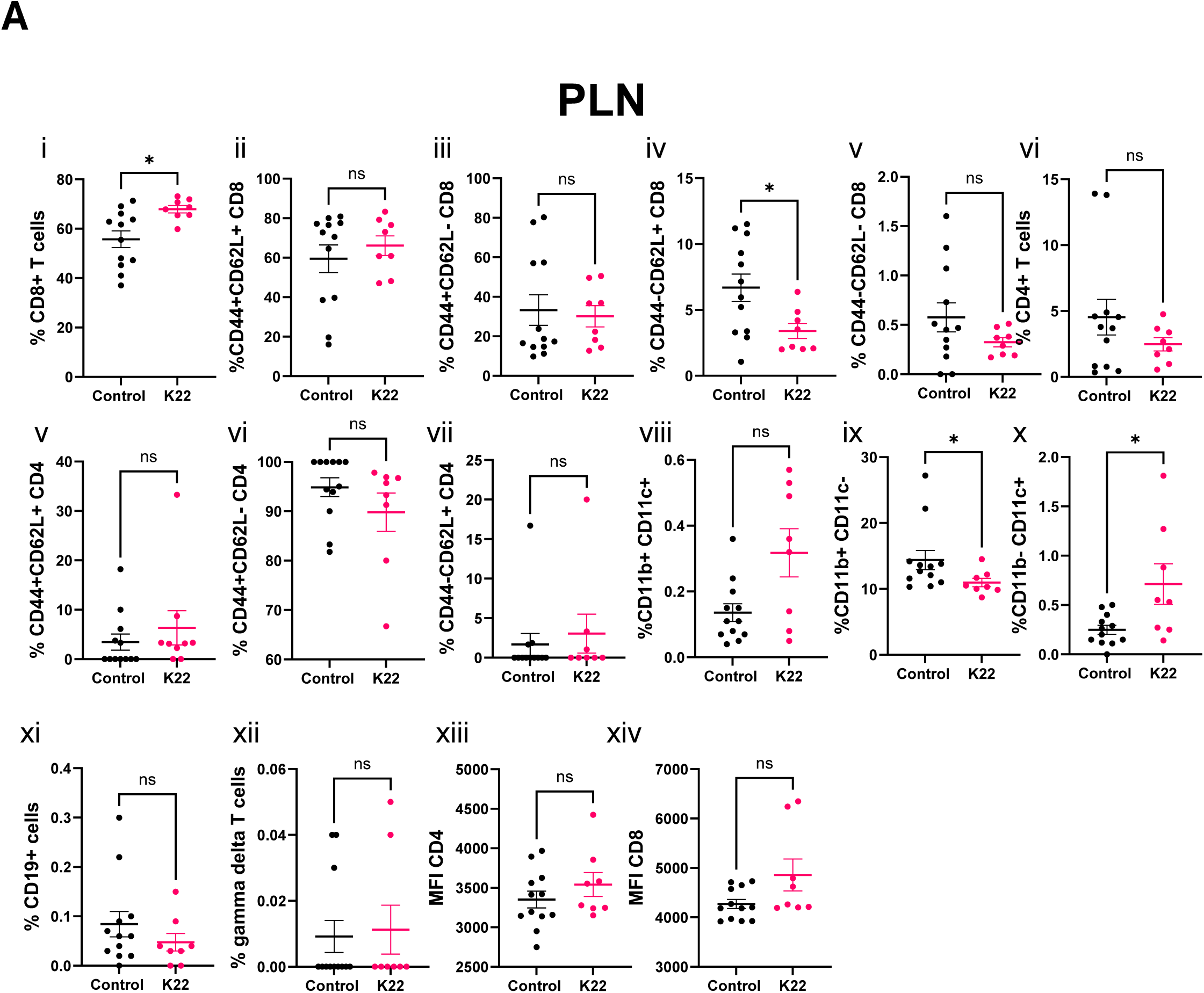
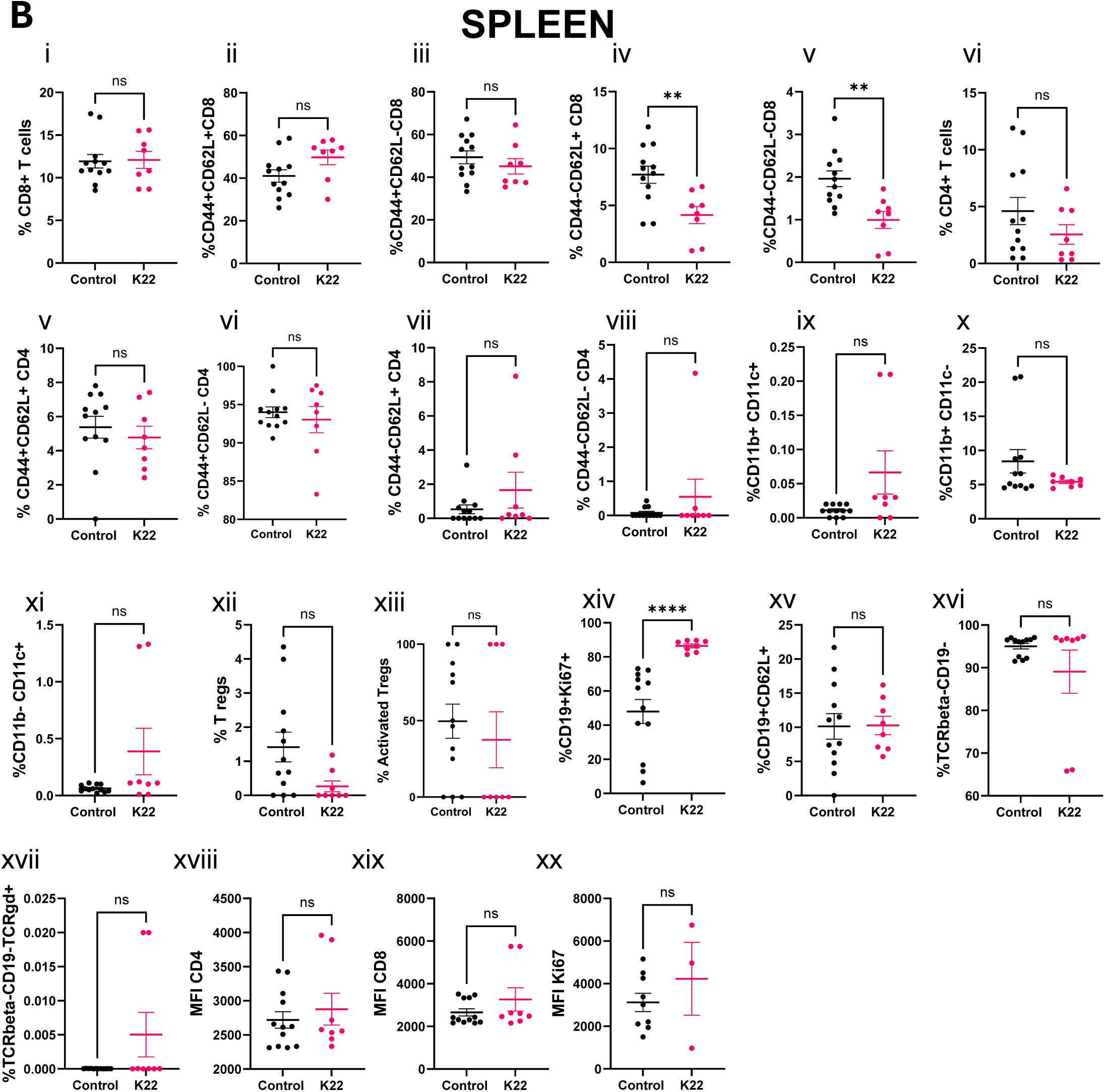
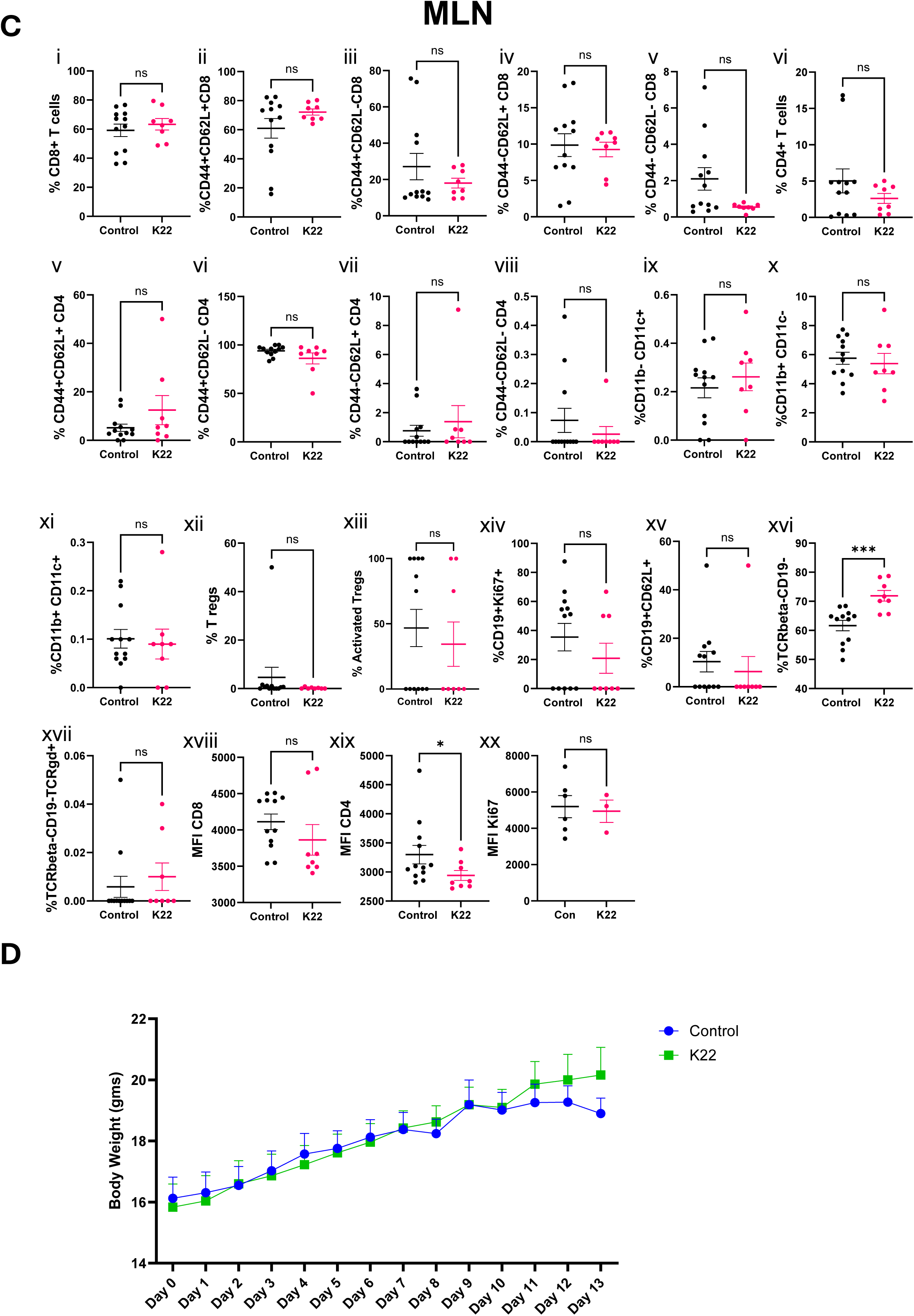
| K22 induces modest, compartment-specific changes in peripheral lymphoid and myeloid subsets and does not affect body weight in the NY8.3 Rag1⁻/⁻ model. Detailed flow-cytometric analysis of immune subsets in PLN (A), spleen (B) and MLN (C) from control and K22-treated mice at endpoint. Panels depict frequencies of total CD8⁺ T cells, naïve and effector CD8⁺ T cells (CD44/CD62L subsets), total CD4⁺ T cells and CD44/CD62L CD4⁺ subsets, CD11b⁺CD11c⁺ and CD11b⁺CD11c⁻ myeloid cells, CD19⁺ B cells (including CD19⁺CD62L⁺ and CD19⁺Ki67⁺ subsets), TCRβ⁻CD19⁻TCRγδ⁺ γδ T cells, and Foxp3⁺ regulatory T cells where available, as well as mean fluorescence intensity (MFI) of CD4, CD8 and Ki67 on T cells. Each symbol represents an individual mouse, bars denote mean ± s.e.m., and statistical comparisons between control and K22 groups (ns, not significant) indicate modest reductions in naïve CD8⁺ T cells and selected myeloid subsets in draining lymph nodes, with no evidence of broad systemic immune dysregulation. (D) Body weight recorded daily from day 0 to day 13 shows no significant difference between control and K22-treated mice, indicating that K22 was well tolerated over the treatment period.

